# Alcam-a and Pdgfr-α are essential for the development of sclerotome derived stromal cells that support hematopoiesis in vivo

**DOI:** 10.1101/2022.03.02.482709

**Authors:** Emi Murayama, Catherine Vivier, Anne Schmidt, Anne-Lou Touret, Philippe Herbomel

## Abstract

Mesenchymal stromal cells are essential components of hematopoietic stem and progenitor cell (HSPC) niches, regulating HSPC proliferation and fate decisions. Their developmental origins are largely unknown. In zebrafish, we previously found that the stromal cells of the caudal hematopoietic tissue (CHT), a niche functionally homologous to the fetal liver in mammals, arise from the ventral part of caudal somites. We have now discovered that this ventral domain is actually the sclerotome, and that two typical markers of mammalian mesenchymal stem/stromal cells, Alcam and Pdgfr-*α*, are distinctively expressed there and instrumental for the emergence and migration of stromal cell progenitors, which in turn conditions the proper assembly of the vascular component of the CHT niche. Furthermore, we find that the trunk somites are similarly dependent on Alcam and Pdgfr-*α* to produce mesenchymal stromal cells that foster the initial emergence of HSPCs from the dorsal aorta. Thus the sclerotome contributes essential stromal cells for each of the key steps of developmental hematopoiesis, and likely is the embryological origin of most if not all mesenchymal stem/stromal cells found in non-cephalic tissues.

## Introduction

Hematopoietic stem and progenitor cells (HSPCs) are multipotent precursors that have self-renewal capacity and continuously replenish all mature blood cells throughout the life span. In zebrafish as in mammalian development, they initially emerge from the dorsal aorta of the embryo through an endothelial hematopoietic transition (EHT)^1,2,3,4^. Then they enter the bloodstream to reach their first niche, the caudal hematopoietic tissue (CHT)^5^, where they expand and undergo their first multi-lineage differentiation^6^. Thus the CHT in fish is the hematopoietic homolog of the fetal liver in mammals. In our previous work, we showed that the CHT niche was mainly composed of a transient venous plexus arisen from the primitive caudal vein, and stromal reticular cells (SRCs) interconnecting as well as lining the branches of this venous plexus^5,7^.

We then discovered that the progenitors of these stromal cells initially arose from the ventral part of the caudal somites through an epithelial mesenchymal transition (EMT)^7^. These stromal cell progenitors (SCPs) then emigrated as strings of 2-3 cells further ventrally, in close interaction with the endothelial cells sprouting from the primitive caudal vein that would make up the venous plexus^7^. This early interaction of the precursors of the two cell types that would constitute the CHT niche was presumably important for the very structure and functionality of that niche. In the present study we have more closely characterized the process of SCP emergence and emigration from the caudal somites and identified important molecular actors in this process. Moreover, since the sclerotome compartment of the somites lies at their ventro-medial side, and undergoes EMT and subsequent emigration to produce the various sclerotome derivatives^8,9^, we have addressed its relationship with the ventral cell clusters of the caudal somites that give rise to the SCPs.

Activated leukocyte cell adhesion molecule (Alcam or CD166) is a large cell surface glycoprotein of the immunoglobulin superfamily that mediates adhesion through homophilic interactions in various tissues^10^, and heterophilic interactions with CD6 at the interface of T cells and antigen-presenting cells^11^, and with galectin-8 in the extracellular matrix^12^. While expressed in a wide variety of tissues, Alcam is often found in cell subsets involved in dynamic growth and/or migration processes including neural development^13^, pharyngeal pouch formation^14^, angiogenesis^15^, kidney development^16^ hematopoiesis^17^, and tumor progression^18^; it is also found in mesenchymal stem cells (MSCs) / bone marrow and fetal liver stromal cells^19,20,21^. The short cytoplasmic tail of Alcam was shown to interact indirectly with the actin cytoskeleton via actin-binding proteins such as ezrin and syntenin-1^22^. These interactions of the cytoplasmic domain dynamically regulate the clustering of Alcam molecules at the cell surface and the strength of the homophilic and heterophilic interactions^23,24^.

Platelet-derived growth factor receptor α (PDGFR-α) is a highly conserved receptor tyrosine kinase. Binding of PDGFs to PDGF receptors induces their dimerization, which unlocks their tyrosine kinase activity and results in autophosphorylation of specific tyrosine residues, which then act as binding sites for intracellular Src homology 2 (SH2) domain-containing signaling molecules^25,26^. PDGFR-α mediated signaling regulates various processes of embryonic development and organogenesis, notably the proliferation then migration and differentiation of specialized mesenchymal cells in various organ anlages^27^. Like Alcam, PDGFR-α is also a marker of bone marrow MSCs^19,28^.

Here we identify Alcama and Pdgfr-*α* as two essential transmembrane proteins in the process of SCP emergence from somite epithelial cells, their subsequent migration to become stromal cells, and for the resulting structure and functionality of the CHT niche. We further show that the somite ventral clusters giving rise to them coincide with the caudal sclerotome, and that Alcama and Pdgfra are similarly important for the emergence from the trunk somites of sub-aortic mesenchymal cells that are essential for the induction of HSPCs from the aorta.

## Results

### Alcama is required for stromal cell development from the somites

We previously described that cell clusters containing SCPs appeared at the ventral side of caudal somites by 21-22 hpf^7^. Somite formation occurs in rostro-caudal direction and we found that the SCP clusters first become apparent by live VE-DIC microscopy in the transition from somite maturation stage S4 to S5 (4th and 5th somite from the tailbud)^29^ (Fig. 1A). During cluster formation, the cells located on the ventral side of the somites adopt a rosette-like structure with a small cavity in the center (Fig. 1A-a; Movie 1). To visualize the subsequent emigration of SCPs from these ventral clusters (VCs), we previously used TCF-driven fluorescent reporters expressed in all somite cells, which were merely inherited by the emigrating SCPs^7^. To analyze the fate of these SCPs, we first used the *Tg(pax3a:eGFP)* and *Tg(ET37:eGFP)* lines^30^. *Tg(pax3a:eGFP)* labelled somite cells weakly, then the VCs more strongly, as well as the migrating SCPs up to 2.5 dpf (Movie 2). The *Tg(ET37:eGFP)* line also highlights somite cells weakly^30^, but then starts to label SCPs more strongly from the onset of their emigration from the somites (Fig. S1A). While these two lines proved useful for our study, we also set out to create a new Tg line that would label the SCPs more specifically and throughout their development. We firstly searched for genes that seemed specifically expressed in the VCs in the ZFIN whole-mount in situ hybridization (WISH) databank. Our subsequent confirmation by WISH of candidate genes between 23-48 hpf led us to target the chondroitin sulfate proteoglycan 4 (*cspg4*) gene (Fig. S1B). We then generated a BAC-based Tg line expressing the GAL4 transcription factor (TF) from the *cspg4* regulatory regions. This *TgBAC(cspg4:GAL4)* line crossed with a *Tg(UAS:RFP)* reporter line faithfully recapitulates the *cspg4* gene expression pattern spatiotemporally (Figs. 1B, S1C). With the exception of the notochord, RFP expression highlights only the VCs and their derivatives in *Tg(cspg4:GAL4;UAS:RFP)* embryos. By 29 hpf, the GFP signal from *Tg(ET37:eGFP)* was also clearly visible and the migration of RFP^+^GFP^+^ cells was observed towards not only the ventral but also the dorsal side (Fig. S1C). The latter cell population migrated along the medial face of the somites towards the notochord (Fig. S1D), suggesting chondro-progenitor and/or tenocyte fates^8,9,31^. Ventral-wards migrating SCPs gave rise to stromal cells throughout the CHT by 38 hpf, and some migrated even further ventrally to become fin mesenchymal cells^30^ (FMCs) in the caudal fin (Fig. 1B-b). These three fates of somite VC derived cells are recapitulated in Fig. S1D.

**Figure 1.**
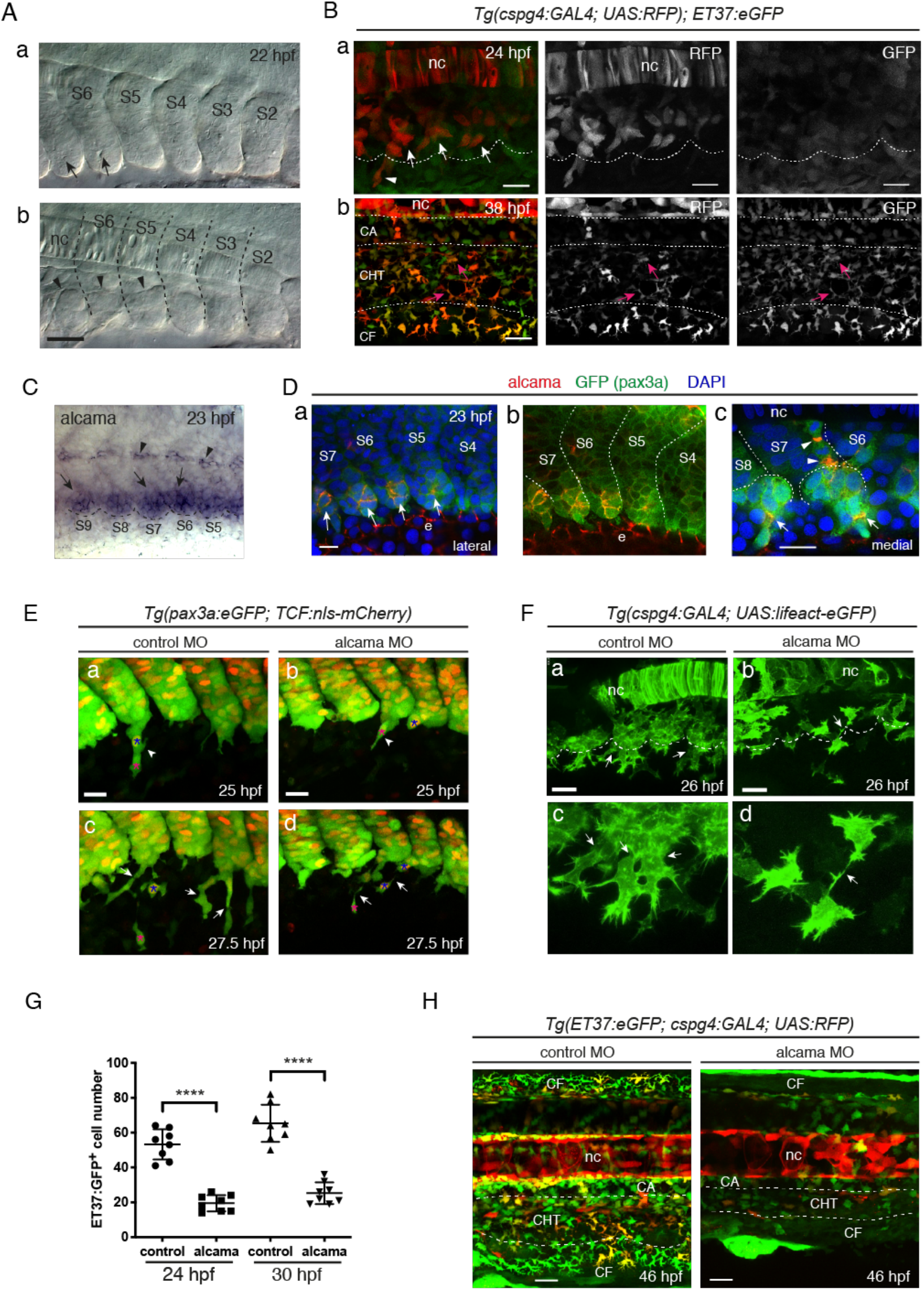
SCP development and requirement for Alcama. **A**, VE-DIC microscopy of developing caudal somites at 22 hpf, rostral to the left, dorsal to the top. Somites are numbered S2-S6 according to their maturation stage counted from the tailbud. Lateral (**a**) and more medial (**b**) focal planes show the central lumen (**a**, arrows) formed within the ventral somite clusters, and the dorsal border of the clusters (**b**, arrowheads). Dashed lines indicate intersomitic boundaries. Scale bar, 40 µm. **B**, Confocal projections of live *Tg(cspg4:GAL4; UAS:RFP; ET37:eGFP)* embryos at 24 (**a**) and 38 (**b**) hpf. **a**, Arrows and arrowhead point at VC cells and an emerging SCP, respectively. The dotted line outlines the ventral border of caudal somites. Magenta arrows in **b** point at RFP/GFP double-positive cells in the CHT. Scale bars, 20 µm. **C**, WISH for alcama at 23 hpf. Arrows and arrowheads point at somite VCs and cells prefiguring the future horizontal myoseptum, respectively. A dotted line outlines the somite ventral border. **D**, Immunostaining for Alcama (red), pax3a:eGFP (green), together with DAPI (blue) at 23 hpf. **a**, Overlay of the red, blue and green signals; arrows point at Alcama labeling in the somite VCs. **b**, Same image as **a** without DAPI staining. e, epidermis. **c**, At a deeper focal plane, strong Alcama signals were detected at contacts between cells migrating semi-collectively dorsal-wards (arrowheads) and ventral-wards (arrows). Dashed lines delineate the VCs and somite borders. Scale bars, 20 µm. **E**, Confocal maximum projections of control or alcama MO injected *Tg(pax3a:eGFP; TCF:nls-mCherry)* embryos at 25 (**a**,**b**) and 27.5 (**c**,**d**) hpf. (**a**,**b**) Arrowheads show SCPs in the early stages of migration; magenta and blue asterisks point to the same cells as in panels **c** and **d**, respectively. (**c**,**d**) Arrows point at the intercellular junctions of migrating SCPs, which are much thinner in **d**. Two blue asterisks in **d** indicate that the similarly labelled cell in **b** has undergone mitosis. Scale bars, 25 µm. **F**, Confocal projections of control or alcama MO injected *Tg(cspg4:GAL4; UAS:Lifeact-eGFP)* at 26 hpf. Dashed lines indicate the somite ventral borders. Arrows point at the connections between migrating SCPs. Magnified images are shown in **c** and **d**. Scale bars, 20 µm. **G**, Quantification of ET37:eGFP^+^ cells in the ventral tail of live control and alcama MO injected embryos, at 24 and 30 hpf over a 5-somites width (n=8 embryos each; mean±SD; ****, P<0.0001. Student’s *t*-test). At 24 hpf, all GFP^+^ cells ventral to the notochord were counted; at 30 hpf, all GFP^+^ cells ventral to the somites were counted. **H**, Confocal projections of control or alcama MO injected *Tg(ET37:eGFP; cspg4:GAL4; UAS:RFP)* embryos at 46 hpf. Dashed lines delineate the CHT. nc, notochord; CA, caudal artery; CHT, caudal hematopoietic tissue; CF, caudal fin.

To understand how somite epithelial cells compartmentalize to give rise to the VCs and their derivatives, we also searched for cell adhesion molecules expressed specifically in the VCs. We found that *alcama* (activated leukocyte cell adhesion molecule a, aka CD166), was expressed in the VCs more strongly than in the bulk of the somites at 23 hpf (Fig. 1C). Immunofluorescence allowed us to detect the Alcama protein at the very core of the clusters as they first appeared morphologically, i.e. at somite stage S5 (Fig. 1D-a,b). Upon somite maturation, Alcama protein signals then gradually propagated to delineate cell boundaries within the clusters (Fig. 1D-a,b). We then detected the Alcama protein at joints between cluster-derived cells migrating both ventral- and dorsal-wards (Fig. 1D-c). We first examined the impact of alcama knock-down by using an antisense morpholino oligonucleotide (MO), that completely blocked translation of the *alcama* transcript in vivo (Fig. S1E,F). Injection in *Tg(pax3a:eGFP;TCF:nls-mCherry)* embryos revealed that while in control embryos, SCPs initially migrated as strings of 2-3 cells interconnected by a rather large contact surface (Fig. 1E-a,c), which immunostaining showed to be enriched in Alcama protein (Fig. 1D-c), in alcama morphants, adhesion between migrating SCPs was reduced to a very small surface at the tip of an elongated cell protrusion (Fig. 1E-b,d). We next crossed our *TgBAC(cspg4:GAL4;UAS:RFP)* line with *Tg(UAS:Lifeact-eGFP)*^32^, in which Lifeact-eGFP allows to highlight F-actin in vivo, so as to visualize actin dynamics and cellular projections during SCP emergence and migration. Lifeact-eGFP^+^ VC-derived cells were clearly less numerous in alcama morphants by 26 hpf (Fig. 1F) - a finding confirmed in the *Tg(ET37:eGFP)* background (Fig. 1G). This reduced number was later mirrored by a similarly (2.7-fold) lower number of SCPs after migration into the CHT by 30 hpf (Fig. 1G), and a clearly lower number of both stromal cells in the CHT and FMCs by 46 hpf (Fig. 1H).

### Alcama regulates the migration behavior of SCPs by modulating F-actin

Time-lapse imaging of migrating Lifeact-GFP^+^ SCPs in alcama morphants revealed a characteristic cell morphology compared to SCPs in control embryos (Fig. 2A). Filopodial projections in migrating leader SCPs usually occur mainly at the leading edge, whereas they were 50% more numerous (Fig. S2A-a) and more randomly scattered around the migrating cell in alcama morphants (Fig. 2B). In addition, even though they were slightly longer on average in alcama morphants (Fig. S2A-b), unlike in control embryos their length did not correlate with their position relative to the direction of migration (Fig. 2C). A further quantitative analysis of confocal live images showed that i) a high Lifeact-GFP (F-actin) signal was present throughout migrating SCPs in alcama morphants whereas the F-actin signal was observed predominantly at the leading edge in controls (Fig. S2B), and (ii) the surface area of migrating SCPs in alcama morphants was much larger than in controls (Fig. S2C). These changes in morphology and F-actin distribution correlated with a strong defect in SCP migration, as their linear distance of migration in alcama morphants was less than half that in the control group (Fig. 2D). Later on, by 46 hpf, this migration defect had translated in a much thinner distribution of ET37:GFP^+^ stromal cells in the CHT (Fig. 1H).

**Figure 2.**
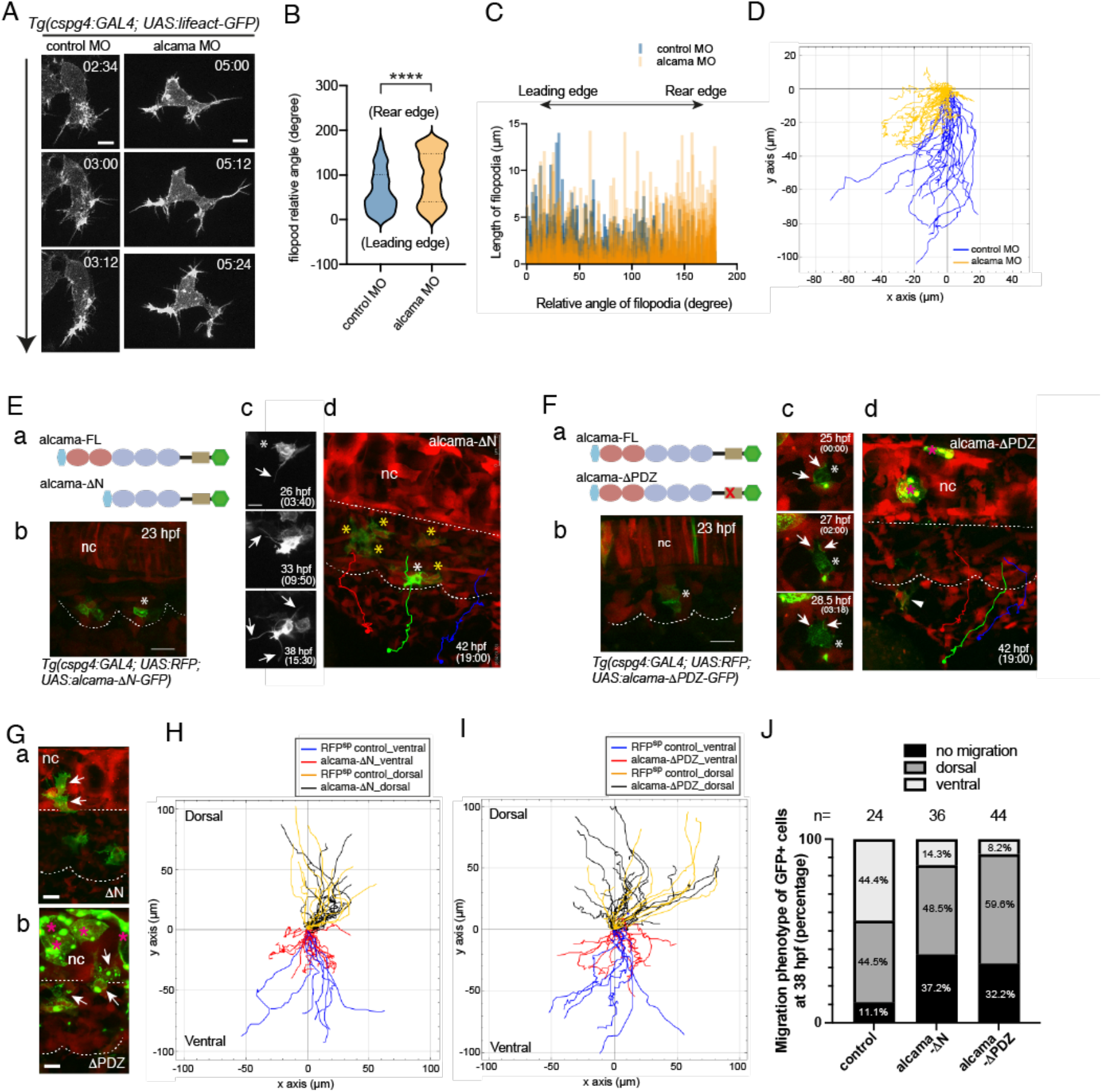
Alcama regulates SCP migration. **A**, Time-lapse confocal images of a migrating SCP leader cell from control or alcama MO injected *Tg(cspg4:Gal4; UAS:Lifeact-GFP)* embryos around 28 hpf. The time point from the initiation of SCP migration is shown in the upper right corner of each panel. The time lag between control and alcama morphant SCPs is due to the fact that the latter tend to take more time to fully egress from the somite. Arrow indicates the direction of cell migration. Scale bars, 10 µm. **B**, Violin plot of filopodia angles relative to the direction of migration for leader SCPs in control and alcama morphant embryos (n=7 SCPs for each group, from 3 independent experiments; median±SD; ****, *P*<0.0001; Mann-Whitney test). **C**, Filopodia length as a function of their relative angles for leader SCPs in control and alcama morphants. **D**, Overlay of individual tracks of control and alcama-deficient SCPs during their migration towards the CHT. In a trajectory plot, all (x, y) coordinates of the SCPs’ starting points (in the clusters) are set to (0,0) and negative or positive values on the Y axis indicate ventral-wards or dorsal-wards migration, respectively. n_Exp_=4, n_embryos_=6, n_tracked cells_=19 and 17 for control or alcama morphants, respectively. **E, a**, Schematic representation of Alcama-FL-eGFP and Alcama-*Δ*N-eGFP mutant constructs. Pink and blue ovals indicate the V-type and C2 type Ig-like domains, respectively. The beige rectangle represents the short cytoplasmic domain. Light blue and green hexagons represent the signal peptide and GFP, respectively. **b**, Appearance of Alcama-ΔN-eGFP^+^ SCPs at 23 hpf. White asterisk indicates the same cell as shown in **c** and **d**. Dotted lines delineate somite and notochord borders. Scale bar, 20 µm. **c**, Morphological evolution over 15.5 hrs of the Alcama-ΔN-eGFP^+^ SCP shown in **b** (See also Movie 4). Scale bar, 10 µm. **d**, End point of the time-lapse imaging shown in (**b**,**c)** and Movie 4. GFP^+^ SCPs marked by yellow asterisks did not migrate out of the somites, whereas RFP^+^/GFP^-^ WT SCPs showed a normal emigration pattern exemplified by the three colored trajectories. A straight dotted line indicates the notochord lower border. **F, a**, Schematic representation of Alcama-FL-eGFP and Alcama-ΔPDZ-eGFP mutant constructs. The red X in the beige rectangle indicates the mutated PDZ binding motif in the cytoplasmic domain. **b**, Appearance at 23 hpf of an Alcama-ΔPDZ-eGFP expressing VC cell (asterisk) followed up over time in **c** (see also Movie 5). Scale bar, 20 µm. **c**, Arrows point at large contact areas between the GFP^+^ cell and neighboring RFP^+^ cells. **d**, End point of the time-lapse imaging of the same embryo shown in **b**; the GFP^+^ SCP followed up in (**b**,**c)** had disappeared by 42 hpf, while another such cell displayed short-distance migration (arrowhead). RFP^+^/GFP^-^ (WT) SCPs showed a normal emigration pattern indicated by three colored trajectories. Straight and curvy dashes lines delineate the ventral border of notochord and somites. **G**, Dorsal-wards migration of Alcama-ΔN-eGFP and -ΔPDZ-eGFP expressing cells (arrows) by 27 hpf. Scale bars, 10 µm. Magenta asterisks in **F**-**d** and **G**-**b** label GFP^+^ notochord cells. **H**, Overlay of individual tracks of control (RFP^+^/GFP^-^ SCPs; orange and blue lines) and Alcama-ΔN-eGFP expressing SCPs (black and red lines). n_Exp_=5, n_embryos_=7, n_tracked cells_=29 and 17 for Alcama-ΔN-eGFP^+^ and control RFP^+^/GFP^-^ SCPs, respectively. **I**, Overlay of individual tracks of control (RFP^+^/GFP^-^ SCPs; orange and blue lines) and Alcama-ΔPDZ-eGFP^+^ SCPs (black and red lines). n_Exp_=4, n_embryos_=5, n_tracked cells_=32 and 17 for Alcama-ΔPDZ-eGFP^+^ and control SCPs, respectively. **J**, Frequency histogram of D/V migration patterns of Alcama-ΔN-eGFP^+^ and -ΔPDZ-eGFP^+^ cells compared with GFP^+^ cells of *Tg(cspg4:Gal4;UAS :GFP)* embryos at 38 hpf. The number of embryos analyzed was 24 (WT), 36 (Alcama-ΔN) and 44 (Alcama-ΔPDZ) from three independent experiments. nc, notochord.

To gain more insight into the implication of Alcama in SCP migration, we took advantage of *Tg(cspg4:Gal4)* driver line to express specifically in these cells three different forms of Alcama fused to eGFP at its C-terminus – a WT full-length Alcama (Alcama-FL-eGFP), and two mutant forms, i) Alcama-*Δ*N-eGFP, lacking both N-terminal Ig-like V-type domains that are known to mediate homophilic cell adhesion^33^, and ii) Alcama-*Δ*PDZ-eGFP, in which we introduced in the short cytoplasmic domain of Alcama three amino acid changes predicted to suppress the binding of PDZ domain containing proteins (Fig. S2D), such as syntenin^22^. These different forms were inserted downstream of a UAS promoter, and the constructs were injected in *Tg(cspg4:Gal4;UAS:RFP)* embryos. Expression of Alcama-FL-eGFP in the VCs did not perturb SCP emergence and migration (Fig. S2E, Movie 3). In contrast, SCPs expressing Alcama-*Δ*N-eGFP appeared disconnected from their neighbors within the VCs, and soon developed an intense filopodial dynamics while still within the somite (Fig. 2Ea-c, Movie 4), but most often with no resulting ventral-wards migration towards the CHT, whereas internal control neighboring RFP^+^GFP^-^ cells migrated normally (Fig. 2E-d). Alcama-*Δ*PDZ-eGFP^+^ VC cells and their SCP derivatives displayed much less filopodia compared to WT or Alcama-*Δ*N-eGFP expressing cells; they still showed some GFP enrichment at cell-cell contacts, but no clear polarity (Fig. 2Fa-c, Movie 5). Like Alcama-*Δ*N-eGFP expressing cells, Alcama-*Δ*PDZ-eGFP^+^ SCPs barely migrated to the ventral side, compared with (RFP^+^GFP^-^) SCPs in the same embryo (Fig. 2F-d). However, the dorsal-wards migration was rather retained for GFP^+^ cells in both cases, as well as for Alcama-FL-eGFP expressing cells (Figs. 2G, S2F,G). In vivo tracking of GFP^+^ cells in both Alcama-*Δ*N-eGFP and -*Δ*PDZ-eGFP expressing embryos confirmed that the cells that egressed ventrally from the clusters then wandered around and migrated only shortly towards the ventral side, whereas the dorsal-wards migration was similar to that of wild-type SCPs or Alcama-FL-eGFP expressing cells (Fig. 2H,I). Statistical analysis of their fates confirmed that the phenotype of GFP^+^ mutant cells was predominantly ‘immobile’ or ‘dorsal migration’, with occasional ‘ventral migration’ limited to a short distance (Fig. 2J). Altogether, these data show that Alcama is cell-autonomously involved in SCP emergence and migration.

### Alcama and PDGFR-*α* deficiency similarly affect SCP development

A prominent feature of alcama morphant and dominant-negative mutant phenotypes was the inhibition of SCP ventral migration. Therefore, we investigated the gene expression of receptors for cytokines and growth factors that might be involved in SCP migration. We found that *pdgfr-α*, a bone marrow stromal cell marker gene in mammals^28^, was specifically expressed in the somite VCs, starting by somite maturation stage S6, and then in the migrating cells derived from them (Fig. 3F-a,c). Therefore we hypothesized a possible cross-talk between Alcama and Pdgfr-*α*. It is known that PDGF signaling leads to an increase in AKT and ERK phosphorylation^34,35^. We firstly investigated the impact of *alcama* knock-down on pERK signaling in the caudal region at 26 hpf. The number of pERK^+^ cells was drastically decreased in the CHT of alcama morphants, while it recovered to almost the control level in the rescued group (Fig. 3A). Then, we analyzed the impact of Pdgfr-*α* deficiency, using a previously validated MO against *pdgfra*^36^. First, we analyzed the colocalization of pERK-positive cells with SCPs marked by pax3a-GFP in alcama and pdgfra morphants. While in control embryos we could observe pERK^+^GFP^+^ migrating SCPs, in alcama and pdgfra morphants the overall number of pERK^+^ cells was reduced, and no pERK signal was observed among migrating SCPs (Fig. S3A).

**Figure 3.**
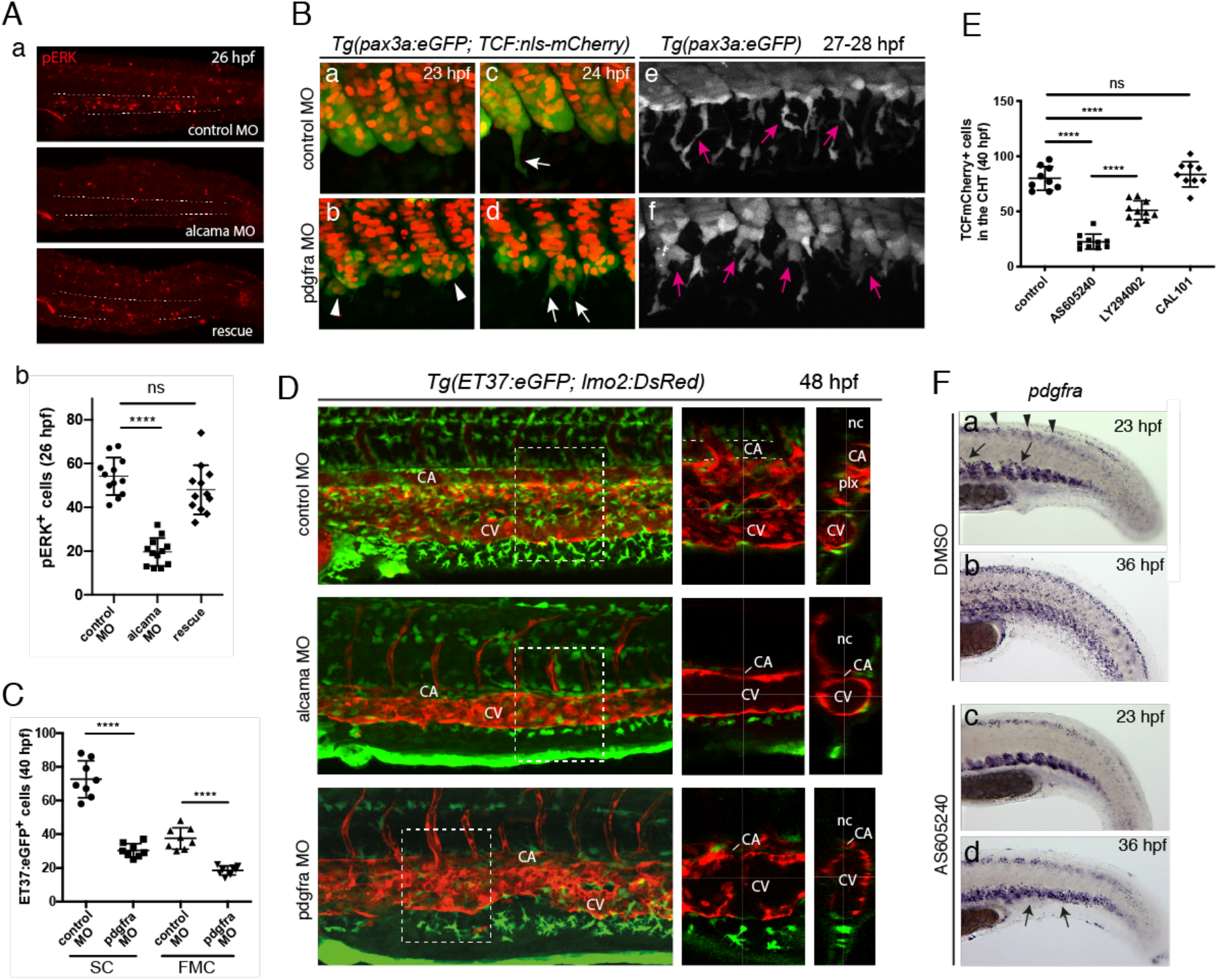
PDGFR*α*/PI3K signaling regulates SCP migration. **A, a**, Immunofluorescence for pERK at 26 hpf in the tail region of embryos injected with control MO, alcama MO, or alcama MO + alcama mRNA (“ rescue”). Representatives of 12 embryos from two independent experiments are shown. Dashed lines indicate the border of CHT. **b**, Quantification of **a** (n=12 embryos for each group; mean±SD; ****, *P*<0.0001, Student’s *t*-test). **B**, Confocal projections of live control (**a**,**c**,**e**) or pdgfra (**b**,**d**,**f**) MO injected *Tg(ET37:eGFP; TCF:nls-mCherry)* embryos from 23 to 28 hpf. Arrowheads point at cohesion defects observed in the VCs of pdgfra MO injected embryos. White and magenta arrows indicate emerging (**c**,**d**) and migrating (**e**,**f**) SCPs, respectively. **C**, Quantification of ET37:eGFP^+^ stromal cells and FMCs in control and pdgfra MO injected embryos at 40 hpf (n=8 for each group, from the same experiment; mean±SD; ****, *P*<0.0001, Student’s *t*-test). **D**, Confocal projections of control, alcama and pdgfra MO injected *Tg(ET37:eGFP; lmo2:DsRed)* embryos at 48 hpf. An enlarged view of the area enclosed by the dashed square is shown in the middle panel, and as an optical transverse section in the right panel. **E**, Quantification of TCF:nmCherry^+^ stromal cells in the CHT of control embryos (n=9), or embryos treated from 20 hpf with AS605240 (n=10), LY294002 (n=10), or CAL101 (n=9) at 40 hpf. Mean±SD; ****, *P*<0.0001, Student’s *t*-test. **F**, WISH for pdgfra at 23 and 36 hpf in 0.2 % DMSO (control) and AS605240-treated embryos. Arrows and arrowheads in **a** indicate VC cells migrating towards the notochord and dorsal mesenchymal cells in the control embryo at 23 hpf, respectively. Arrows in **d** show SCPs slightly dispersed around the VCs at 36 hpf. CA, caudal artery; CV, definitive caudal vein; Plx, venous plexus; nc, notochord.

Time-lapse imaging of SCP emergence in pdgfra morphants by 23-24 hpf revealed that the cohesion of the VCs was compromised, with groups of cells often delaminating together from the somite (Fig. S3B), and then several SCPs per somite cluster or cell group starting to emigrate simultaneously (Fig. 3B), yet with a clearly reduced migration efficiency relative to WT. Quantification showed that the numbers of CHT stromal cells and FMCs originating from SCPs were reduced 2.4-fold and 2.0-fold, respectively, in pdgfra morphants by 40 hpf (Fig. 3C), similar to what we previously found for alcama morphants (Fig. 1H). Live imaging also showed that like in alcama morphants, the extent of ventral-wards migration of SCPs by that time was much reduced in the pdgfra morphants (Movie 6), and this apparently directly impacted on the co-migration of endothelial cells that made the venous plexus, resulting in a much thinner venous plexus, often reduced to a single convoluted tube, in tight apposition to the overlying caudal artery, which could locally show a reduced diameter (Fig. 3D). We then attempted to identify PDGF ligands that would activate Pdgfr-*α* signaling in the VCs and migrating SCPs. pERK immunostaining was performed on embryos overexpressing either of three different PDGFs whose ventro-caudal specific expression had been confirmed by WISH (Fig. S3C). All three ligands were found to activate pERK signaling, hence likely Pdgfr-*α* signaling; however the effect of Pdgfaa was statistically most significant (Fig. S3D).

### PDGFR-*α* drives SCP migration through PI3K signaling

PI3K signaling is known to be activated downstream of various growth factor receptors including PDGF receptors^37^ following their activation. Pharmacological inhibition of PI3K isoforms using AS605240 (inhibiting PI3K isoforms γ, α), LY294002 (inhibiting isoforms α, β, δ) or CAL101 (inhibiting isoform δ) revealed that the migration of SCPs was compromised in embryos treated with LY294002, and more so with AS605240, resulting in a 3.5-fold reduction in stromal cell numbers by 40 hpf (Fig. 3E), whereas pax3a-GFP^high^ neural crest cells migrated with only a slight delay (Fig. S3E, blue arrows). To monitor SCP migration over longer intervals, AS-treated embryos were fixed and labeled with a pdgfra in situ probe at stages encompassing the entire SCP migration process. In the control group, dorsal-wards migration of VCs derived cells had already started by 23 hpf, whereas no migration was observed in the AS-treated embryos (Fig. 3F-a,c). By 36 hpf, SCP migration had not started yet in the AS-treated group, in which SCPs accumulated at the zone of their emergence, while it was already complete in the control group (Fig. 3F-b,d).

Since the involvement of PI3K in SCP migration was revealed, we then mutated two tyrosine residues in the intracellular domain of Pdgfr-α that are responsible for the binding of class IA (α/ α/ δ) PI3K to Pdgfr-*α* following its activation and autophosphorylation^34^ (Fig. 4A), and we cloned the mutated or WT pdgfra ORF followed by a tandem HA-tag downstream of a hsp70 promoter^38^ (hsp70:pdgfra-ΔPI3K-HA or hsp70:pdgfra^WT^-HA, respectively). Transient transgenics resulting from injection of the hsp70:pdgfra-ΔPI3K-HA or hsp70:pdgfra^WT^-HA construct in *Tg(pax3a:GFP)* embryos were heat-shocked at 20 hpf, then imaged in vivo from 23 hpf, or fixed for immunofluorescence at the same time point. The anti-HA antibody positive signals sometimes showed a mosaic distribution, but were sufficiently present in the targeted tissue of hsp70:pdgfra-ΔPI3K-HA and hsp:70:pdgfra^WT^-HA embryos at 24 hpf (Fig. S4A). Heat-shocked hsp70:pdgfra-ΔPI3K-HA embryos typically showed a less clear outline of the somite VCs than controls. In addition, 38% of them showed multiple SCPs simultaneously initiating migration from one somite, reflecting the lack of leader cell, and 20.5% had at least one somite showing an ‘overflow’ phenotype in which the VC collectively delaminated into the ventro-caudal cavity, reminiscent of the phenotype observed above in pdgfra morphants; these cell populations tended to disperse over time, but did not go through the normal migration process (Figs. 4B, S4B). In embryos overexpressing the pdgfra^WT^-HA construct, more apoptotic bodies were detected than in WT embryos, but no abnormalities were observed in the VCs and emigration of SCPs (Fig. S4C). Interestingly, the morphology of migrating SCPs in pdgfra-ΔPI3K expressing embryos was quite similar to that observed in alcama morphants, with long and numerous filopodia and the cell’s main axis often orthogonal to the ventral-wards direction of migration (Fig. 4C, Movie 7), and with increased 3D cell surface area and volume (Fig. S4D,E). Importantly, the number of VCs derived GFP^+^ cells was clearly more decreased for those migrating ventral-wards (SCPs) than for those migrating dorsal-wards (DMCs) (Fig. 4D), again as previously observed upon Alcama deficiency. The total number of SCPs at 36 hpf in the CHT of pdgfra-ΔPI3K expressing embryos was also strongly reduced (Fig. 4E), as in Alcama- or Pdgfra-deficient embryos.

**Figure 4.**
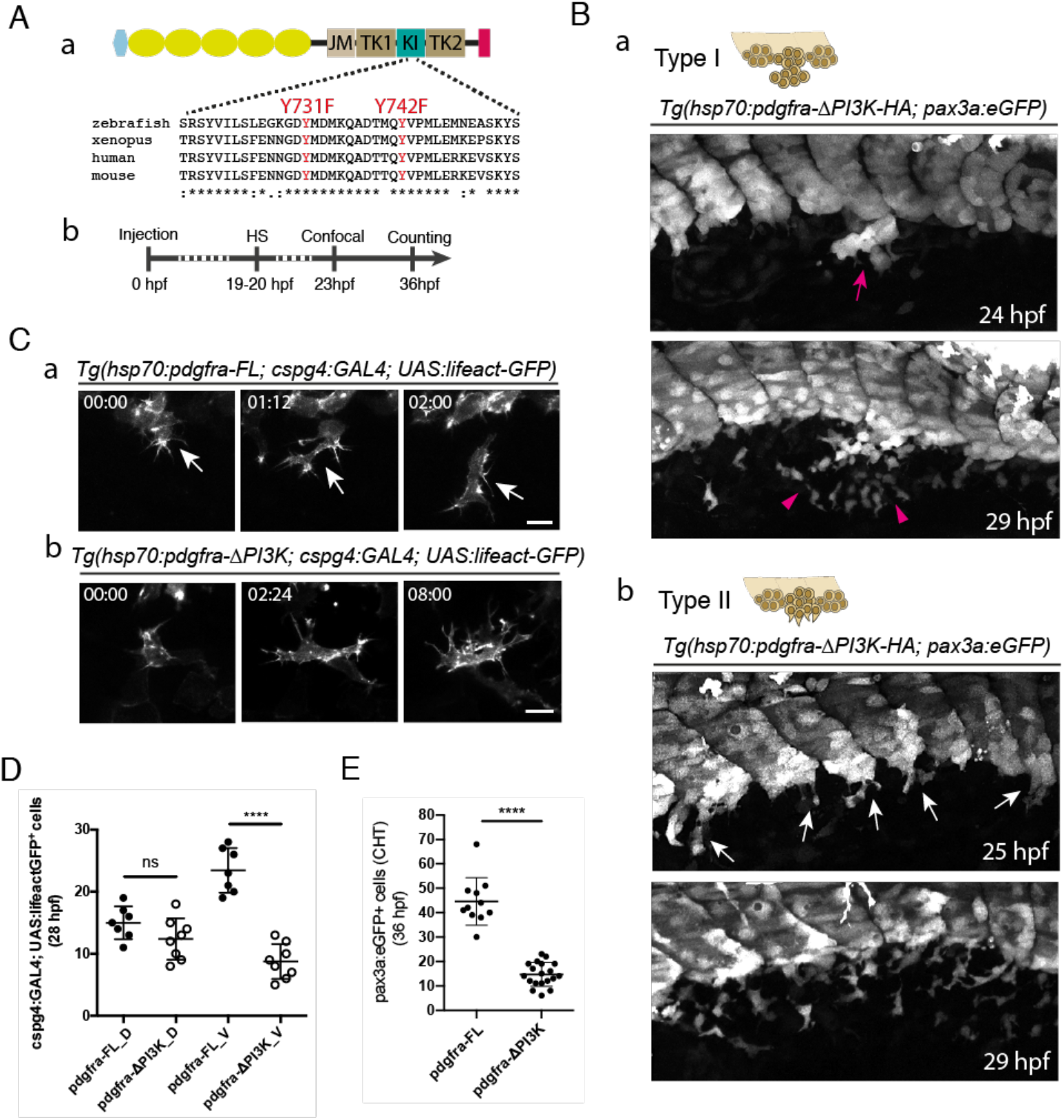
Pdgfra influences cluster cohesion and the following migration behaviour of SCPs through PI3K binding. **A. a**, Schematic representation of Pdgfra fused to tandem HA-tag (red) at its C-terminus, and amino acid alignment around the site of mutations in the KI (kinase insert) domain. The light blue hexagon and yellow ovals indicate the signal peptide and Ig-like extracellular domains, respectively; JM, juxtamembrane; TK1, tyrosine kinase domain 1; TK2, tyrosine kinase domain 2; tyrosine residues involved in PI3K binding when phosphorylated that we mutated into phenylalanines are shown in red. **b**, Time course of the experiment; HS, heat-shock. **B**. Confocal projections from live *Tg(hsp70:pdgfra-ΔPI3K; pax3a:eGFP)* embryos showing Type I (**a**) and Type II (**b**) cluster cohesion / emergence defects (see also Fig. S4B). Magenta arrow in **a** indicates SCPs overflow out of the cluster before the initiation of migration. This cell population was dispersed in situ after 5 hours (magenta arrowheads). **b**, Arrows indicate multiple SCPs emerging simultaneously from the cluster. **C**, Representative frames from a confocal time-lapse sequence of individual migrating SCP leader cell in *Tg(cspg4:GAL4; UAS:lifeact-eGFP)* embryos injected with the hsp70:pdgfra-FL (**a**) or hsp70:pdgfra-*Δ*PI3K (**b**) construct and heat-shocked at 20 hpf. Confocal imaging was performed from 23 hpf. Time is indicated in hrs and min. Scale bar, 25 µm. **D**. Quantification of *Tg(cspg4:GAL4; UAS:lifeact-eGFP*^*+*^) SCPs migrating dorsal-wards (D) or ventral-wards (V) in Pdgfra-FL (control) and Pdgfra-*Δ*PI3K expressing embryos at 28 hpf (mean±SD; n=7 embryos for control, n=8 for hsp70:pdgfra-*Δ*PI3K, from 3 independent experiments. ****, *P*<0.0001, Student’s *t*-test. **E**, Counting of pax3a:eGFP^+^ SCPs in control and hsp70:pdgfra-*Δ*PI3K embryos in the CHT at 36 hpf, over a 5-somite width (mean±SD; n=11 for control, n=19 for hsp70:pdgfra-*Δ*PI3K, from 3 independent experiments. ****, *P*<0.0001, Student’s *t*-test.

### Alcama and Pdgfr-*α* expression are regulated by sclerotomal transcription factors

Since our live imaging revealed that the somite VCs giving rise to SCPs also gave rise to cells migrating dorsal-wards towards the notochord, it suggested an overlap or identity of SCP clusters with the sclerotome. Our WISH analysis revealed that TFs considered as sclerotome markers, such as Pax9, Snai2, Twist1a and Twist1b were indeed expressed in the VCs of caudal somites from the S4/S5 somite stage, and also quite earlier - from the S1 stage - for Snai2 (Fig. 5A). Therefore, we investigated if these TFs regulate the expression of *alcama* and *pdgfra* during SCP development. To this end, we first isolated promoter regions of the *alcama* (3.3 kb) and *pdgfra* (3.2 kb) genes from BACs, and cloned them into the pGL3 Basic vector to monitor their promoter activity in vivo in various conditions through a Dual luciferase assay performed on tail lysates at 23 hpf (Fig. 5B-a,E-a).

**Figure 5.**
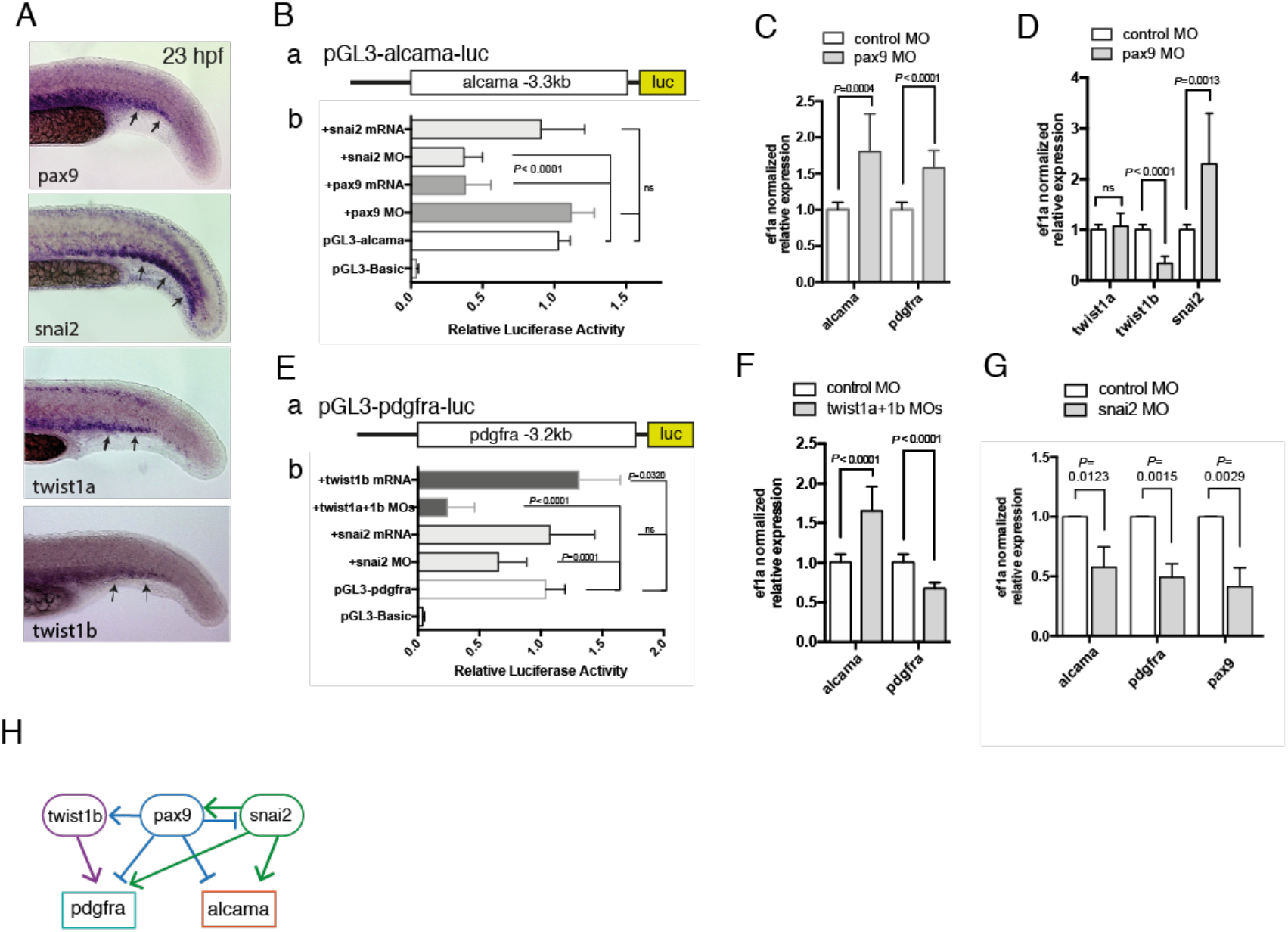
*alcama* and *pdgfra* are regulated by sclerotome-associated TFs. **A**, WISH for *pax9, snail2, twist1a* and *twist1b* at 23 hpf in the tail of WT embryos. Arrows point to expression of these genes in the somitic VCs. **B, a**, a DNA fragment encompassing the 3.3 kb region upstream of the of *alcama* gene transcription start site was inserted upstream of the luciferase (luc) coding region of the pGL3-Basic plasmid, to give pGL3-alcama-luc. **b**, in situ luciferase assays using pGL3-alcama-luc, which was injected at the 1-cell stage either alone or together with the indicated mRNA or MO; lysates obtained from 10-15 tails at 23 hpf for each condition were subjected to luciferase assay (n=9 for each, as biological triplicates from three independent experiments) or pax9 (n=6 for each). The empty vector pGL3-Basic and pGL-alcama-luc (n=11 for each) were used as negative and positive controls, respectively. **C**, qPCR analysis for *alcama* and *pdgfra* in pax9 morphants and control (n=9 for each, a set of biological triplicates from three independent experiments) at 23 hpf. **D**, qPCR analysis for *twist1a, twist1b* and *snai2* in pax9 morphants and control (n=9 for each) at 23 hpf. **E**,**a** a DNA fragment covering the 3.2 kb region upstream of the *pdgfra* gene transcription start site was inserted upstream of the luciferase (luc) coding region of the pGL3-Basic plasmid, to give pGL3-pdgfra-luc. **b**, Same experimental scheme as in B-b, following injection of the indicated constructs and Mo or mRNA combinations. **F**, qPCR analysis for *alcama* and *pdgfra* (n=6 for each) in twist1a/twist1b double morphants and control (n=9) at 23 hpf. **G**, qPCR analysis for *alcama, pdgfra* and *pax9* in snai2 morphants and control (n=3) at 23 hpf (mean±SD). **H**, A proposed network for the regulation of TFs acting on alcama and pdgfra during SCP development. Line ending with arrow or bar represent activation or repression, respectively.

Co-injection of the pGL3-alcama-luc construct with either snai2 MO or pax9 mRNA attenuated the alcama promoter activity, while co-injection with snai2 mRNA or pax9 MO had no significant effect (Fig. 5B-b). Consistent with this, alcama mRNA expression in the tail of pax9 morphants was upregulated at 24 hpf (Fig. 5C), while it was down-regulated in the tail of snai2 morphants (Fig. 5G). We similarly analyzed the effects of *twist1a/twist1b* and *snai2* on the *pdgfra* promoter by co-injection with pGL3-pdgfra-luc. Double knockdown of *twist1a* and *1b* led to a drastic decrease in *pdgfra* promoter activity in the tail samples, whereas co-injection of twist1b mRNA stimulated it (Fig. 5E-b). In line with this, *pdgfra* mRNA expression in the tail was downregulated in twist1a/1b morphants (Fig. 5F). Snai2 MO also reduced the *pdgfra* promoter activity (Fig. 5E-b), and consistently, pdgfra expression in the tail of snai2 morphants was significantly reduced (Fig. 5G). Finally, we found that these sclerotomal TFs regulated each other in the tail, as *pax9* knockdown up-regulated *snai2* expression but down-regulated *twist1b*, while *snai2* knockdown down-regulated *pax9* (Fig. 5D,G). The network of genetic interactions deduced from all these data is depicted in Fig. 5H.

### Alcama and Pdgfr-*α* are also essential for the production of mesenchymal derivatives of trunk somites, and of definitive HSPCs

The expression of Alcama, Pdgfra, and the sclerotomal TFs mentioned above in the ventral-most part of zebrafish somites is not restricted to the tail, but also extends to the somites of the trunk (Fig. 6A). Moreover, as in the tail, pax3a:eGFP^+^ or ET37:eGFP^+^ mesenchymal cells are present in the trunk by 36 hpf, notably just lateral or ventral to the dorsal aorta, in close contact with it or with the dorsal wall of the underlying axial vein (Fig. 6B,6C-a,6D-a, and Movie 8). As in the caudal region, we found that Alcama, Pdgfr-a or Pax9 deficiency led to a strong reduction of these pax3a:GFP^+^ or ET37:eGFP^+^ stromal cells of the trunk (Fig. 6C,D). In addition, these cells were found lateral or ventro-lateral to the aorta, but never just ventral to it (Fig. 6C,D). By 52 hpf, the total number of GFP^+^ stromal cells in the trunk of all morphants was even more reduced relative to control embryos than at 36 hpf (Fig. S5).

**Figure 6.**
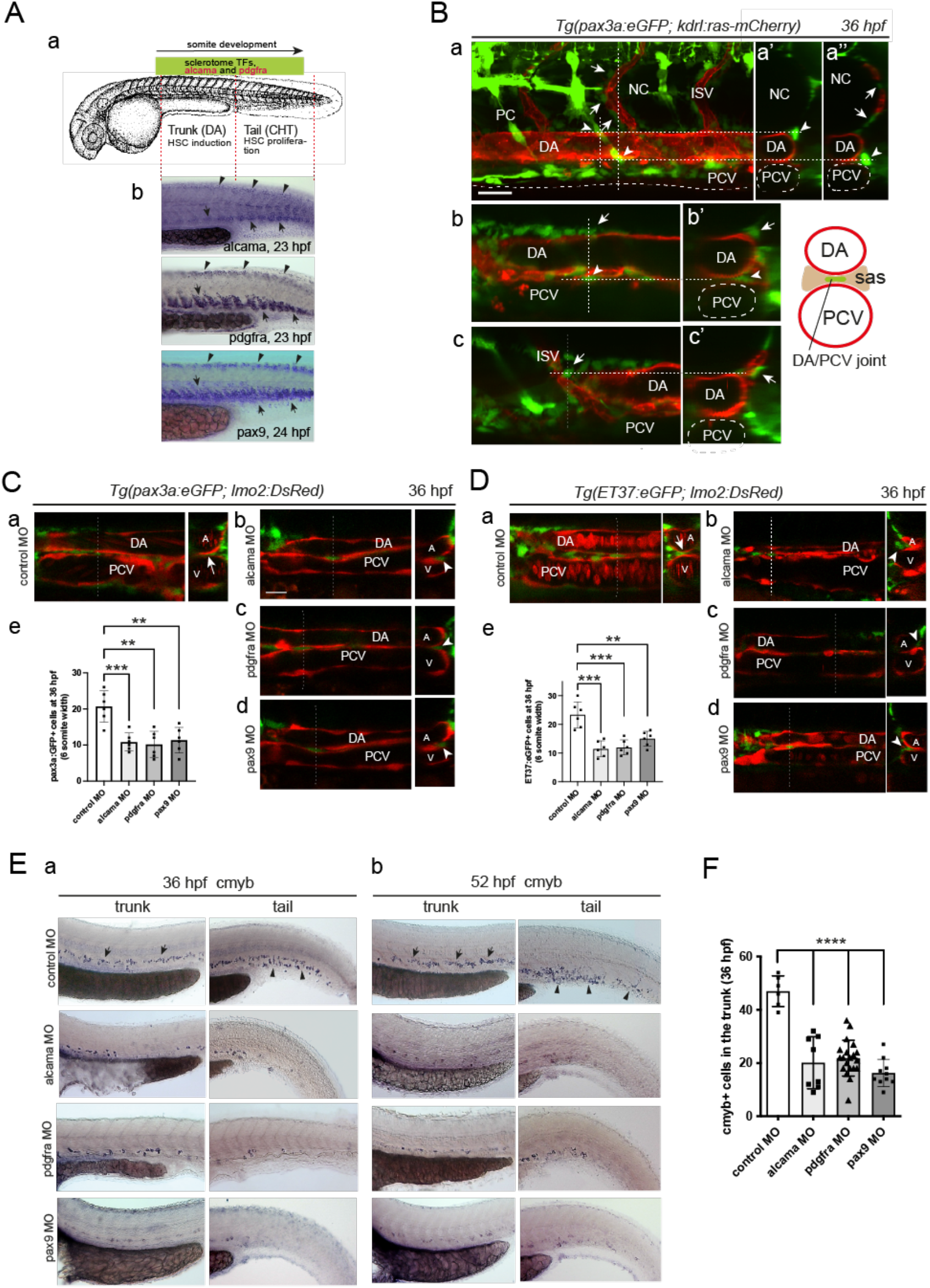
Alcama and Pdgfra are also involved in stromal cell development in the trunk, which conditions the development of definitive HSPCs. **A, a**, rostro-caudal extent of sclerotomal TFs and *alcama* and *pdgfra* expression by 24 hpf. **b**, WISH for *alcama, pdgfra* and *pax9* at 23-24 hpf. Arrows point at their expression in the somite VCs/sclerotome in the trunk and tail. Arrowheads point at their expression in the smaller dorsal sclerotome^31^. **B**, Confocal maximum projection (**a**) and single confocal sections (**b**,**c**) of the trunk region of a *Tg(pax3a:eGFP; kdrl:ras-mCherry)* embryo at 36 hpf, with vertical dashed lines showing the position of the optical transverse sections shown on the right (**a’**,**a’’**,**b’**,**c’**). Arrowheads point at GFP^+^ stromal cells located at the DA/notochord or DA/PCV junction, arrows point at ISV-associated mesenchymal cells, known to originate from the dorsal-wards migration of ventral sclerotome cells^31^,^70^. Scale bar, 20 µm. (**C**,**D**) Confocal sections of the trunk region of *Tg(pax3a:eGFP; lmo2:DsRed)* (**C**) and *Tg(ET37:eGFP; lmo2:DsRed)* (**D**) embryos injected with control (**a**), alcama (**b**), pdgfra (**c**) or pax9 (**d**) MOs. Dashed lines indicate the position of the corresponding transverse section shown on the right. Arrows and arrowheads point at stromal cells located at the DA/PCV joint or somewhat lateral to it, respectively. **e**, Quantification of pax3a:eGFP^+^ (**C**) or ET37:eGFP^+^ (**D**) cells in the sub-aortic space of 36 hpf live embryos injected with control, alcama, pdgfra or pax9 MO (mean±SD; n=6 embryos for each group. ***, *P ≤* 0.001; **, *P ≤* 0.01; Student’s *t*-test). Counting was done over a 6-somites width. **E**, WISH for *myb* at 36 (**a**) and 52 (**b**) hpf in embryos injected with control, alcama, pdgfra or pax9 MO. Arrows and arrowheads point at myb^+^ signals in the trunk (arrows) and tail (arrowheads) regions, respectively. **F**, Quantification of myb^+^ cells at 36 hpf as shown in **E**-**a** (mean±SD; n=6 embryos for control, 8 for alcama, 21 for pdgfra and 10 for pax9 MO injected embryos. ****, *P≤*0.0001, Student’s *t*-test). DA or A, dorsal aorta; PCV or V, posterior cardinal vein; ISV, intersomitic vessel; sas, sub-aortic space.

36 hpf is the developmental stage by which definitive HSPCs start to emerge by EHT from the ventral wall of the aorta - a process that peaks by 48-52 hpf. We found that Alcama, Pdgfr-*α* or Pax9 deficiency all caused a dramatic reduction in myb^+^ HSPCs in the trunk by 36 hpf (Fig. 6E,F). At the same time-point, we could observe the beginning of myb^+^ HSPC colonization of the CHT of control embryos, whereas almost no myb^+^ cell was detected in the CHT of all morphants (Fig. 6E-a). Then, at 52 hpf, only few myb^+^ HSPCs were found in the trunk of all morphants, even less than at 36 hpf, while myb^+^ cells were numerous in control embryos, and very few myb^+^ HSPCs were detected in the CHT of all morphants (Fig. 6E-b). Thus the severe impact of Alcama or Pdgfr-*α* deficiency on the development of sclerotome-derived stromal cells both in the trunk and in the tail led to a profoundly defective definitive hematopoiesis.

## Discussion

Mouse studies have shown the presence of various stromal cell subsets in hematopoietic stem cell niches^39^. However, the developmental process and behavior of these cells, and the molecules involved in cell-cell interactions during niche formation, remain a mystery. In the present study, we have taken advantage of the accessibility to in vivo observations of the zebrafish CHT, the hematopoietic homolog of the fetal liver niche in mammals, to provide new insights into the genesis of stromal cells. We previously found that the stromal cells of the CHT arose from ventral clusters (VCs) within the caudal somites. We report here a new transgenic line, *TgBAC(cspg4:GAL4*), that specifically labels these VCs and their derivatives, notably the SCPs throughout their development, and at least two other types of mesenchymal cells - dorsal-wards migrating cells (DMCs) and fin mesenchymal cells (FMCs)^30^. Considering that the cell population originating from the caudal somites that differentiates into osteoblasts in adult fish^40^ may also be included, cells derived from the VCs may have the potential of mesenchymal stem cells (MSCs). VC cells specifically express sclerotome marker genes such as *twist1, snai2* or *pax9*, and their anatomical location also supports their sclerotomal nature. The three transmembrane molecules that we found here to be more specifically expressed in the VCs and their derivatives, Cspg4, Alcama and Pdgfr-*α*, have all been used as markers of bone marrow and/or fetal liver MSCs in mammals^19,28,39,41^. Our tracing of the SCPs back to the VCs of the caudal somites allows us to connect the concept of MSCs, which arose from studies of in vitro cell cultures from postnatal mammals, to the embryological concept of sclerotome. The VC-derived DMCs would notably give rise to chondrocytes and tenocytes^8,9,31^. It will be important to study whether the ‘commitment’ process^42^, which corresponds to the first branching point in the differentiation of MSCs into various cell lineages, may already take place within the VCs.

It is the outstanding optical clarity of the ventro-caudal region of zebrafish embryos as the caudal somites develop that allowed us to identify for the first time the earliest morphological individualization of the sclerotome, as a distinct ventral cell cluster appearing at somite maturation stage S5, before these cells become mesenchymal cells migrating both dorsal- and ventral-wards. Interestingly, Naganathan et al.^43^ recently found that up to stage S4, somites undergo a mechanical adjustment of their A-P length that is facilitated by somite surface tension, which requires the somite to be fully packed within an uninterrupted basal lamina. It therefore makes sense that cell migrations out of the somite, which require to break the basal lamina, are “ allowed” to begin only once this mechanical rearrangement of the somite has been completed, i.e. from somite stage S4/S5. Morphological individualization of VCs by stage S5 appears to closely follow the onset of expression at the same place of the sclerotomal TF genes as well as *alcama*. It may thus reflect the EMT triggered by Twist1/Snai2/Pax9. At this stage, Alcama first appears in the center of the cluster and then gradually spreads to all interfaces among cluster cells. Then as these cells emigrate from the cluster, they do so as strings of cells still enriched in Alcama at their interfaces. Suppression of Alcama function reduced the number of emigrating SCPs, their contacts among each other, and their migratory capacity. The latter defect correlated with a less efficient polarization of the cell and its F-actin dynamics relative to the ventral-wards direction of migration. The augmented filopodial dynamics observed in Alcama-*Δ*N expressing SCPs may be favored by the lack of Alcama-mediated cell-cell contacts, while the still present short cytoplasmic domain of this overexpressed Alcama-*Δ*N form may trigger more F-actin dynamics generating filopodia. Conversely, VC cells overexpressing a mutant Alcama form in which the cytoplasmic domain is no longer able to bind PDZ domain containing actin-binding proteins such as syntenin-1^22,23,24^ showed a lack of filopodial dynamics, more extended cell-cell contacts, and an even more complete lack of apparent cell polarity.

Suppression of Pdgfr-*α* function affected the emergence and ventral-wards migration of sclerotome derivatives at least as strongly as Alcama deficiency. Our finding that both Alcama and Pdgfr-*α* deficiencies led to a strong decrease in pERK signaling in the ventro-caudal region suggests that the intracellular signaling pathways downstream of Alcama and Pdgfr-*α* overlap within the stromal derivatives of the sclerotome. In addition, we found PI3K activation to be an obligatory step downstream of Pdgfr-*α* activation for the development of these cells. Interestingly, the dorsal-wards migration of sclerotome cells was also affected in embryos treated with a PI3Kα,γ inhibitor, indicating that even though distinct pathways are likely involved in the dorsal and ventral migrations of sclerotome derivatives, both use PI3K as a necessary downstream effector (Fig. S6).

Clements & Traver^44^ suggested that cells derived from the trunk sclerotome were required for the emergence of HSPCs. Nguyen et al.^45^ then found that somite-derived pax3a:eGFP^+^ cells fostered HSPC emergence, but interpreted them as dermomyotome-derived cells integrating into the DA endothelium. Here we have confirmed their finding but clarified that these pax3a:eGFP^+^ cells are actually sclerotome-derived, DA-associated mesenchymal cells. At least some of them, lateral and ventral to the aorta, are most likely precursors of the mural cells later ensheathing the DA that were shown to arise from trunk somites^46,47^ – more precisely the sclerotome, as in mammals and birds^48,49^, and to initially associate with the aorta floor. Those seen in close association with the dorsal wall of the axial vein (PCV) underlying the aorta are highly reminiscent of the stromal cells that we previously found by electron microscopy bridging gaps between adjacent venous endothelial cells at this very location^6^, which is where aorta-derived HSPCs enter circulation in zebrafish. Altogether our present study leads to the unifying conclusion that the somitic VCs identified as the sclerotome successively give rise to mesenchymal cells involved in the first two main stages of HSPC development and function. Trunk somites first contribute mesenchymal stromal cells that foster the emergence of HSPCs from the neighboring DA floor, then the later formed caudal somites contribute mesenchymal stromal cells that are key components of the first niche where the aorta-derived HSPCs settle, expand and undergo multi-lineage differentiation.

Our study further revealed that Alcama and Pdgfr-*α*, known as mere markers of MSCs in mammals, are actually instrumental for the very emergence and emigration of mesenchymal stromal cells from the somites to build the hematopoietic niches both in the trunk and tail. Our new *Tg(cspg4:Gal4)* line, which highlights mesenchymal stromal cells from their sclerotomal origin onwards, will be a precious tool to discern, notably through single-cell transcriptome analysis, whether cellular heterogeneities prefiguring different fates are already present within the somitic VCs, and how different the stromal cells fostering HSPC emergence from the aorta are from those that then nurture HSPC expansion and multi-lineage differentiation in the CHT niche. Finally, beyond stem/stromal cells found in hematopoietic organs, cells with MSC capacities have been extracted from a variety of fetal and adult mammalian tissues, but their precise origin within these tissues has remained elusive. Crisan et al.^50^ presented data suggesting that these various tissue MSCs were perivascular mesenchymal cells/pericytes. Interestingly, *cspg4* expression, which we found here to mark the early sclerotome and its derived cells, is a typical marker of pericytes^51,52^. Therefore altogether our study makes it reasonable to anticipate that developmentally, MSCs from most if not all tissues posterior to the head will prove to arise from the sclerotome. Elucidation of the molecular mechanisms of MSC development will contribute to the field of regenerative medicine by facilitating the induction and manipulation of rare MSC populations in adult tissues.

## Materials and Methods

### Zebrafish

Fish were maintained in our zebrafish facility at Institut Pasteur. Embryos were obtained through natural crosses, raised at 28°C in embryo water [Volvic® water containing 0.28 mg/ml Methylene Blue (M-4159; Sigma) and 0.03mg/ml 1-phenyl-2-thiourea (P-7629; Sigma)], and staged according to Kimmel et al.^53^ For this study, we used the previously described transgenic lines *Tg(ET37:GFP)*^54^, *Tg(pax3a:EGFP)*^*il150*, 55^, *Tg(UAS:Lifeact-GFP)*^*mu271*, 32^, *Tg(7xTCF-Xla*.*Siam:nlsmCherry)*^56^, *Tg(kdrl:ras-mCherry)*^57^ and *Tg(lmo2:DsRed)*^58^.

### BAC recombineering and transgenesis

The tol2 and BAC recombineering vectors were kindly provided by K. Kawakami (National Institute of Genetics, Mishima). A BAC DNA containing the *cspg4* gene (DKEY-105I23) was purchased from Source BioScience. The *TgBAC(cspg4:GAL4)* line was firstly generated by recombining a iTol2-amp cassette to the BAC vector^59^, then a second recombineering was performed targeting a cassette containing GAL4 (pGAL4FF-FRT-Kan-FRT) flanked by 50 bp arms homologous to the region around the translation start of the cspg4 gene. Primers used for the 1st and 2nd recombineering steps are listed in the Supplementary information (Table 1). Recombination was performed using RedET methodology (K001, GeneBridges) as described in a previous report^60^. BAC DNAs were prepared using NucleoBond BAC 100 (740579, Marchery-Nagel). *Tg(UAS:RFP)* embryos were injected with 1 nl of mix solution of 100 ng/µl BAC DNA and 50 ng/µl Tol2 mRNA at 1-cell stage.

### In situ hybridization, immunostaining and TUNEL

Whole-mount RNA *in situ* hybridization (WISH) was performed according to Thisse (https://wiki.zfin.org/display/prot/Thisse+Lab+-+In+Situ+Hybridization+Protocol+-+2010+update). Riboprobes were synthesized from PCR fragments amplified from cDNA extracted and synthesized from the tails of wild-type embryos as a template, or by reverse transcription of linearized plasmids. The probes used in this study are indicated in the Supplementary information (Table 2a,2b).

Whole-mount fluorescent immunostaining was carried out as described^5^. The list of primary and secondary antibodies and concentrations used is shown in the Supplementary material (Table 3). TUNEL staining was performed using ApopTag Red In Situ Apoptosis Detection Kit (S7100, MerckMillipore) combined with peroxidase (POD) coupled anti-DIG antibody (1/200, #11207733910, Roche) followed by tyramide-based amplification^61^.

### Construction of mutant form of alcama and pdgfra

#### UAS:alcama-ΔN-eGFP, UAS:alcama-ΔPDZ-eGFP and UAS:alcama-FL-eGFP

Firstly, the alcama ORF was cloned to utilize as a template for the following PCRs (details of cloning are indicated below). For *alcama*-*ΔN* construct, a DNA fragment missing to encode amino acids 25-238 corresponding to the two amino-terminal Ig-like domains of Alcama was amplified. For the *alcama*-*ΔPDZ* construct, 3 amino acid substitutions were introduced in a putative PDZ domain binding motif at positions 533-539 (KTRQGSW->**MV**RQGS**G**) by site-directed mutagenesis. Primers used for the mutagenesis are listed in Supplementary information (Table 4). Control alcama-FL (full length) and mutated alcama fragments were cloned into the Nco I site of UAS:eGFP vector in frame with eGFP using Gibson assembly (#E5510, NEB).

#### hsp70:pdgfra-ΔPI3K-HA and hsp70:pdgfra-FL-HA

A zebrafish pdgfra-*Δ*PI3K was designed based precisely on the human dominant-negative pdgfra, with the substitutions of Y706F (Y731F in human) and Y717F (Y742F)^25^. Fragments containing amino acid substitutions were amplified by PCR using the clone containing full-length wt pdgfra (kindly provided by J. Eberhart, University of Texas at Austin) as a template. A 3 kb fragment covering the ORF was amplified in three fragments and the point-mutations were introduced to the appropriate fragments using primers listed in Supplementary information (Table 4). A tandem HA-tag sequence was added in frame at a C-terminal of pdgfra. Fragments encoding control or mutant pdgfra were cloned into hsp70 vector using Gibson assembly.

### Transient transgenesis

Plasmid DNA for each construct was co-injected with capped mRNA coding for the Tol2 transposase into 1-cell stage embryos at a concentration of 250 and 25 ng/µl, respectively.

### RNA/cDNA synthesis, plasmid construction and injection

Total RNA was extracted with the TRIzol reagent (15596026, Invitrogen) from tails of anesthetized embryos at 23 hpf. Tail total RNA was reverse transcribed into cDNA using a Superscript IV reverse transcriptase (18091050, Invitrogen). For *alcama, pdgfaa, pdgfab, pdgfbb* and *twist1b*, ORFs were amplified by PCR using tail cDNA as a template for the subcloning into the TOPO vector and then reinserted into pExpress1 vector using Gibson assembly. ORFs of *pax9* and *snai2* were amplified by PCR from tail cDNA then cloned directly into pExpress1 vector. Primers used for the cloning are listed in Supplementary information (Table 5). mRNAs were synthesized using mMessage mMachine transcription kit (AM1344, Ambion). Capped mRNAs were injected into 1-cell-stage embryos at the amount of 100 pg.

### Morpholinos and qPCR

Morpholino oligonucleotides (Gene Tools) were injected (0.5-1 nl) into 1-cell-stage embryo at the amount specified; alcama MO^14^ (2 ng), pdgfra MO^36^ (4 ng), pax9 MO^62,63^ (6 ng), snai2 MO^63,64^ (8 ng) and twist1a and twist1b MOs^65^ (2 ng each). MO sequences are shown in the Supplementary information (Table 6). For quantitative real-time PCR (qPCR), total RNA were extracted from 3 independent groups of 40 tails dissected from embryos at 23 hpf to synthesize template cDNAs. The primers used for qPCR are shown in the Supplementary information (Table 7). All qPCR experiments were performed with measurements taken from 3 technical replicates. Fold changes in gene expression were calculated using the 2^-ΔΔCT^ method and normalized to ef1a.

### Heat-shock treatment

Embryos were heat-shocked by placing them in pre-warmed Volvic water for 30 min at 39°C then transferred to 28°C until in vivo observation^38^.

### Promoter cloning and Dual-Luciferase assay

3.3 and 3.2 kb promoter regions of *alcama* and *pdgfra* genes were cloned from BAC DNA (*alcama*: RP71-78P1, BACPAC Resources; *pdgfra*: DKEY-97C6, Source BioScience) and cloned into the pGL3 basic vector (E1751, Promega) linearized with Xho I and Hind III (pGL-alcama-luc and pGL-pdgfra-luc). Primers used for promoter cloning are listed in Supplementary information (Table 8). To assess *alcama* and *pdgfra* promoter activities in response to various TFs, WT embryos were injected at the 1-cell stage with luciferase vectors (pGL-alcama-luc or pGL-pdgfra-luc), control (Renilla) expression vector (pRL-TK; E2241, Promega), and specified amount of Morpholino or appropriate mRNA encoding TF (co-injection condition is listed in Supplementary information; Table 9). Tails were dissected from the injected embryos at 23-24 hpf and lysed with passive lysis buffer (E1941, Promega) for subsequent Dual-Luciferase assay following the manufacturer’s instructions (E1910, Promega).

### Drug treatments

Pharmacological inhibitors were all solubilized in DMSO (Sigma) and appropriate DMSO controls (0.2 %) were used for all experiments. Embryos treated with AS605240 (2 µM; S1410, Selleckchem), LY294002 (20 µM; S1105, Selleckchem) and CAL101 (10 µM; S2226, Selleckchem) from 19-20 hpf embryos then live-images were taken with drugs in agarose and embryo water supplemented with tricaine. For WISH, embryos were treated with drug from 19-20 hpf until the stage of interest, then fixed in 4% methanol-free formaldehyde overnight.

### Live imaging, image processing and quantification

VE-DIC (video-enhanced differential interference contrast) microscopy was performed using a Polyvar2 microscope (Reichert) as described previously^66^. Confocal fluorescence microscopy was performed on Leica SPE and SP8 set-ups and Andor spinning disk set-up as described previously^4,7^.

The image processing was basically performed using Fiji^67^. Measurements of surface area and volume on migrating SCPs were performed using IMARIS software (Bitplane). Cell tracking was performed using Manual tracking plugin (Fiji) and the obtained data were 1) plotted with a Python script developed by S. Rigaud (Image Analysis Hub (IAH) of Institut Pasteur) or 2) imported into the Chemotaxis and Migration Tool plugin^68^ (Fiji). For the analysis of filopodial dynamics, Filopodyan plugin^69^ was used on Fiji. First, we obtained time-lapse movies (6 min intervals, 23-38 hpf) of the migrating lifeact-GFP^+^ SCPs injected with control or alcama MO (7 leader cells were analyzed for each condition). We extracted 7-10 timepoints from the time that the leader cell left its followers and measured the orientation of the filopodia for each timepoint. The cut-off threshold was set to 3 µm.

The direction of SCP migration was obtained from the positional information at the first and last timepoints. The angle of filopodia relative to the direction of SCP migration was calculated with a Python script developed by D. Ershov (IAH, Institut Pasteur).

### Statistical analysis and replication

Statistical analysis was performed using GraphPad Prism 9. For data that followed a normal distribution, statistical significance was assessed by a two-tailed Student’s *t*-test, while Mann-Whitney test was introduced for two groups with unequal variances. Significance levels were set at *p<0.05, **p<0.005, ***p<0.0005 and ****p<0.0001. Data are shown as mean or median±SD. All experiments are a representation of at least three independent experiments unless stated otherwise. For qPCR and luciferase assays, results were collected with three biological replicates.

## Acknowledgements

We thank our fish facility team for zebrafish care, K. Kawakami (National Institute of Genetics, Mishima) for providing us with the BAC recombineering plasmids, J. Eberhart (University of Texas at Austin) for the pdgfra:pCR4 plasmid, B. Appel (University of Colorado) for the cspg4:pJC53.2 plasmid, and A. Kudo (Tokyo Institute of Technology, Nagatsuta) for the pax9:pPICT3 plasmid. We also thank Stephane Rigaud, Dmitry Ershov and Jean-Yves Tinevez (IAH of Institut Pasteur) for implementation of Python scripts for filopodia analysis. We gratefully acknowledge K. McElreavey, D. Houzelstein and C. Eozenou (Institut Pasteur) for their help with Luciferase assays. We are also thankful to N. Wolff and M. Genera (Institut Pasteur) for their help with Western blotting.

## Author contributions

E.M. conceived the project, designed, performed, and analyzed experiments, and wrote the manuscript. C.V. conducted experiments and fish care. A.S. subcloned the hsp70 vector and assisted with helpful discussions. A-L.T. conducted experiments. P.H. analyzed, discussed, and suggested experiments, and wrote the manuscript.

## Competing interests

The authors declare no competing interests.

## Funding

This work was supported by Institut Pasteur, CNRS, and by grants to P.H. from the Fondation pour la Recherche Médicale (#DEQ20160334881), the Fondation ARC pour la Recherche sur le Cancer, and the Agence Nationale de la Recherche Laboratoire d’Excellence Revive (Investissement d’Avenir; ANR-10-LABX-73).

## Notes

### Competing Interest Statement

The authors have declared no competing interest.

