## Supplementary material for "Alcam-a and Pdgfr-α are essential for the development of sclerotome derived stromal cells that support hematopoiesis in vivo": Supp Info

### Fig S1

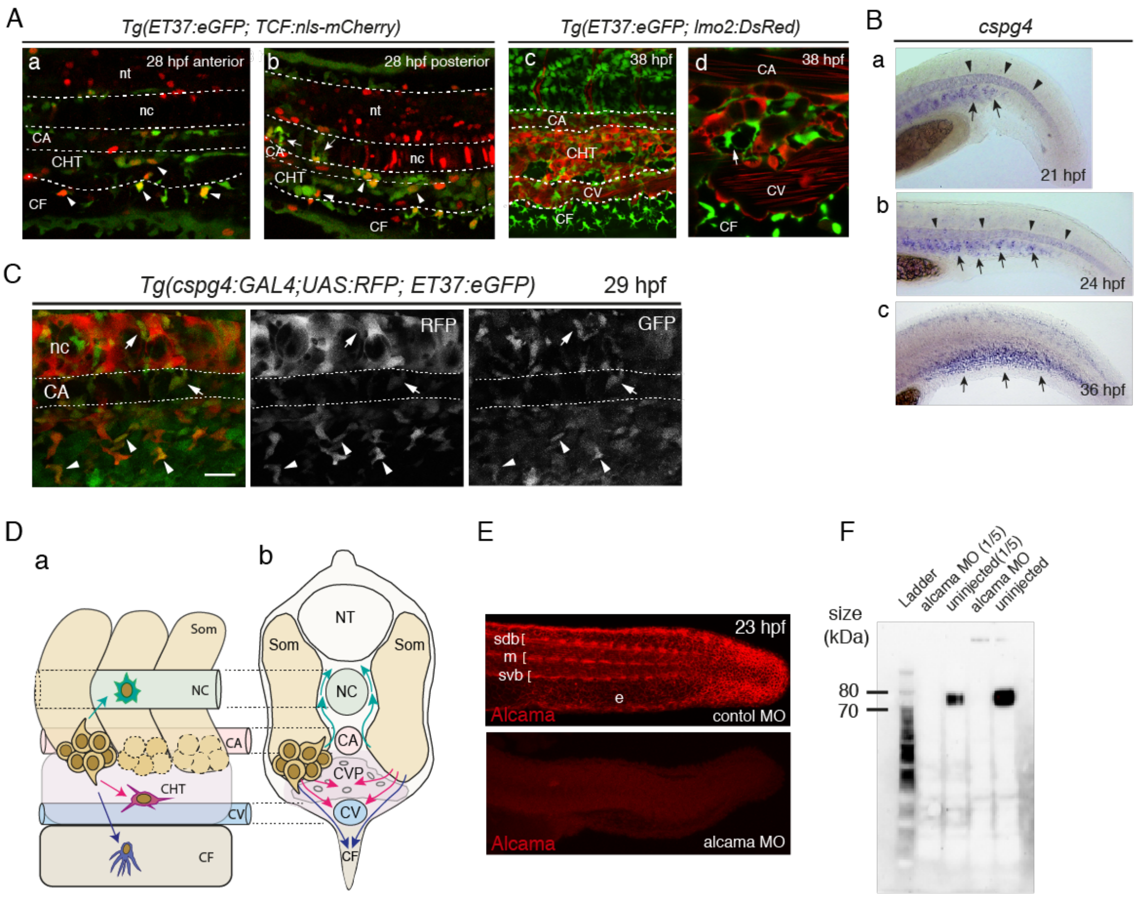


**Figure S1. Transgenic reporter lines used to study SCP development, and effects of alcama knockdown.**

**A.** Confocal projections of the ventro-caudal region of *Tg(ET37:eGFP; TCF:nls-mCherry)* embryos at 28 hpf (**a**,**b**) and *Tg(ET37:eGFP; lmo2:DsRed)* embryos at 38 hpf (**c**,**d**). **B,** WISH for *cspg4* at 21 (**a**), 24 (**b**) and 36 hpf (**c**). Arrows and arrowheads indicate somite VC derived cells and notochord, respectively. **C**, Confocal projection of *Tg(ET37:eGFP; cspg4:GAL4; UAS:RFP)* embryo at 29 hpf. Arrows and arrowheads indicate dorsal- and ventral-wards migrating GFP$^{+}$/RFP$^{+}$ VC derived cells, respectively. Dashed lines delineate the borders of the caudal artery. Scale bar, 20 µm. **D**, Schematic representation of the caudal region in lateral (**a**) and transverse view (**b**), showing the migration paths of somite VC derived cells. Magenta and blue arrows indicate the paths leading to CHT stromal cells and FMCs, respectively. Green arrows show the dorsal-wards migration path (of putative chondrocyte and tenocyte progenitors). **E**, Immunofluorescence for Alcama in control or alcama MO-injected embryo at 23 hpf. **F**, Western blot analysis showing the efficiency of alcama MO using anti-Alcama antibody. Cell lysates were obtained from tails at 24 hpf then whole protein extracts were analyzed at two different dilutions. nt, neural tube; nc notochord; CA, caudal artery; CV, definitive caudal vein; CF, caudal fin; sdb, somite dorsal border; m, slow muscle pioneers prefiguring the horizontal myoseptum; svb, somite ventral border; e, epidermis.

Fig. S2

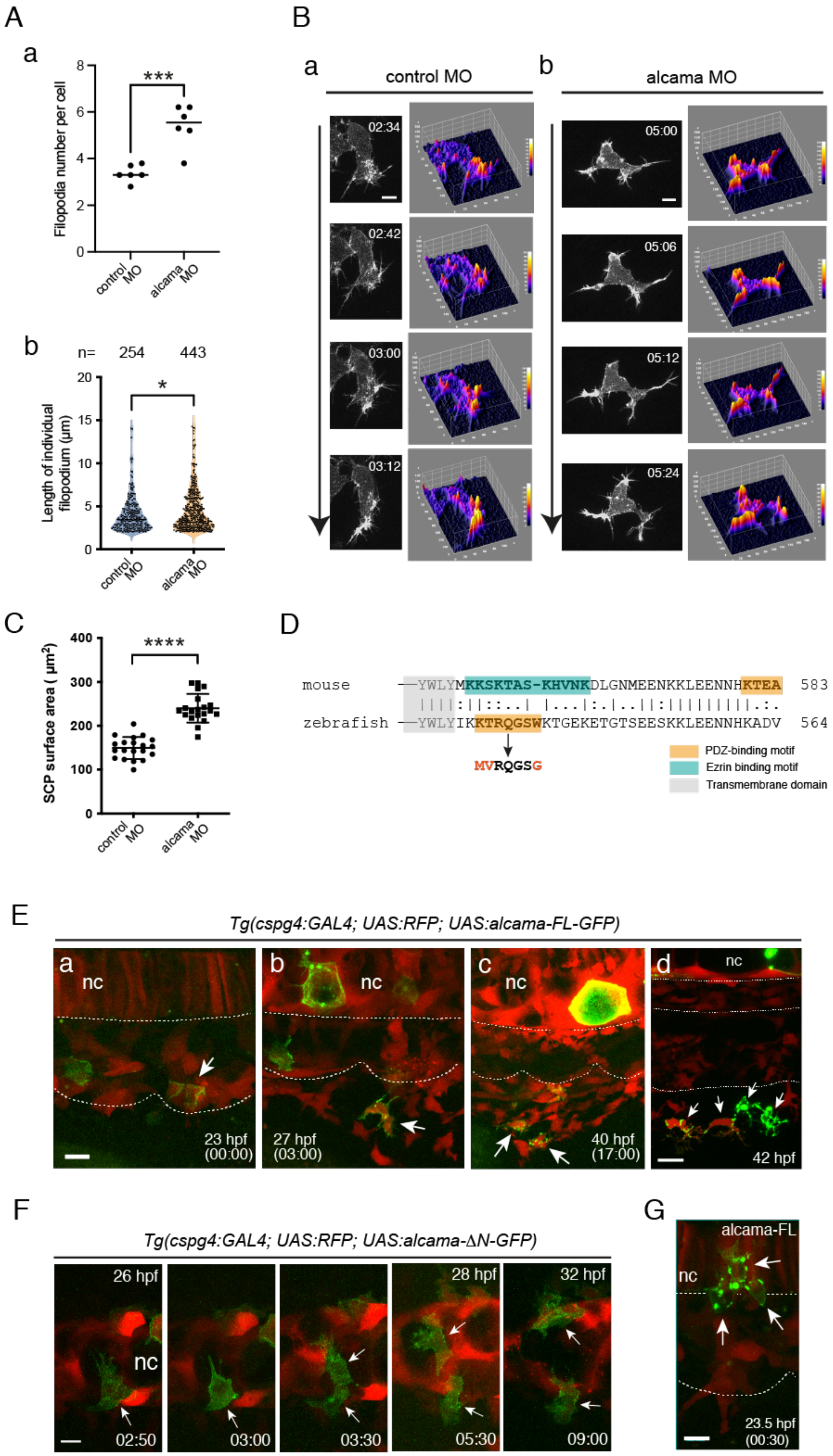

A
a
Filopodia number per cell
***
control MO
alcama MO
b
n= 254 443
Length of individual filopodium (μm)
*
B
a
control MO
02:34
02:42
03:00
03:12
b
alcama MO
05:00
05:06
05:12
05:24
C
SCP surface area ( μm²)
****
D
mouse —YWLYMKKSKTAS-KHVNKDLGNMEENKKLEENNHKTEA 583
zebrafish —YWLYIKKTRQGSWKTGEKETGTSEESKKLEENNHKADV 564
MVRQGSG
PDZ-binding motif
Ezrin binding motif
Transmembrane domain
E
Tg(cspg4:GAL4; UAS:RFP; UAS:alcama-FL-GFP)
a nc 23 hpf (00:00)
b nc 27 hpf (03:00)
c nc 40 hpf (17:00)
d nc 42 hpf
F
Tg(cspg4:GAL4; UAS:RFP; UAS:alcama-ΔN-GFP)
26 hpf
nc
02:50
03:00
03:30
28 hpf
05:30
32 hpf
09:00
G
alcama-FL
nc
23.5 hpf (00:30)

**Figure S2. Alcama modulates SCP migration. A**, Quantification of filopodia number and length for migrating SCP leader cells in control and morphant embryos from the experiments analyzed in Fig. 2A-C (n=7 cells per condition, from 3 independent experiments). **a**, Graph comparing the total number of filopodia ≥ 3 µm per cell during migration (averaged over 6 time-points; mean±SD; *, $P$<0.0168; Student's $t$-test). **b**, Quantification of filopodia length (≥ 3 µm) for migrating SCP leader cells (measured at 7-10 time points; median±SD; *, $P$<0.0189; Mann-Whitney test.). **B**, Surface plots of Lifeact-GFP intensity in the migrating SCPs shown in Fig. 2A; warmer colors represent higher intensity; vertical arrows, direction of migration. **C**, Quantification of 2D surface area of SCPs in the *Tg(cspg4:GAL4; UAS:lifeact-eGFP)* embryos injected with control or alcama MO. Mean±SD; n=20 cells for each, obtained in 8 embryos from three independent experiments. ****, $P$<0.0001; Student's $t$-test. **D**, Alignment of amino acid sequences of the short cytoplasmic domain of mouse Alcam (CD166) and zebrafish Alcama. The light grey shading indicates the last residues of the transmembrane domain. Orange and green shadings indicate PDZ- or Ezrin-binding motif, and the PDZ-binding motif as mutated in this study is shown at the bottom. **E**, Control cells expressing alcama-FL-eGFP construct showed a normal development, exemplified here by a GFP$^+$ cell followed over time from its appearance in a VC by 23 hpf (**a**, arrow), then as a ventral-wards emigrating SCP (**b**, arrow), that later underwent mitosis by 40 hpf (**c**, arrows); scale bar, 10 µm. **d**, Alcama-FL-GFP$^+$ FMCs (arrows) at 42 hpf; scale bar, 20 µm. **F**, Dorsal-wards migration of an Alcama-ΔN-eGFP$^+$ VC cell (arrows) from 26 to 32 hpf; it underwent mitosis by 29.5 hpf, then one of the daughter cells migrated further dorsally. Scale bar, 10 µm. **G,** Dorsal-wards migration of three Alcama-FL-GFP$^+$ VC cells (arrows) at 23.5 hpf. Scale bar, 10 µm. Nc, notochord.

Fig S3

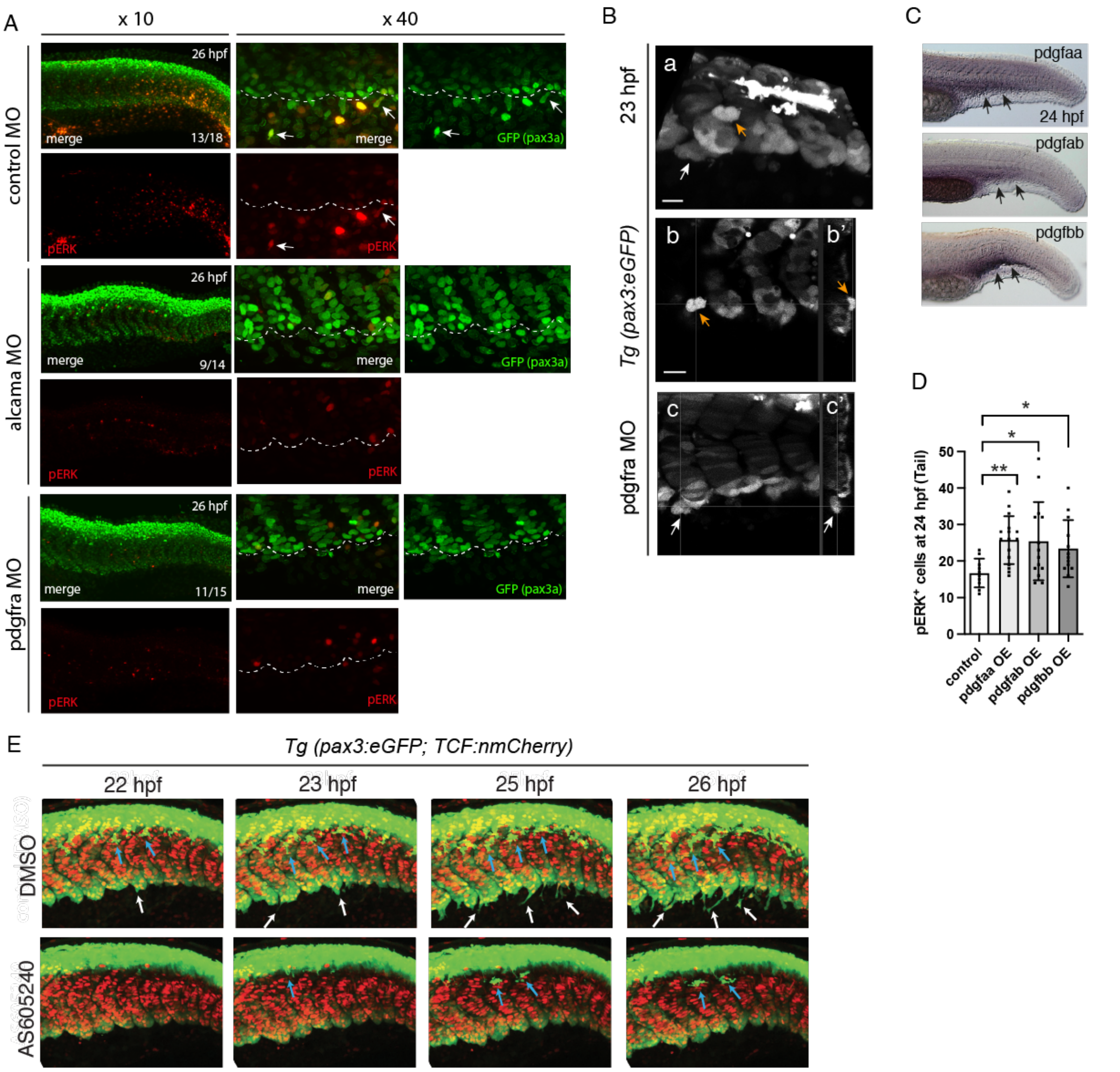


**Figure S3. Involvement of PDGFRα signaling in cluster cohesion and subsequent migration of SCPs. A,** Immunofluorescence at 26 hpf for pERK and GFP in *Tg(pax3a:eGFP)* embryos injected with control, alcama or pdgfra MO. Arrows point at pERK$^+$/GFP$^+$ SCPs and dashed lines delineate the ventral border of caudal somites. **B**, Cluster cohesion defects at 23 hpf in the *Tg(pax3a:eGFP)* pdgfra morphant embryo shown in Fig. 3B-b. White and orange arrows point at cell groups that appear to have detached from a VC laterally and medially. **a**, Ventrally tilted maximum projection view; (**b,c)** single confocal planes; (**b',c')** optical transverse sections at the positions shown by a vertical line and arrow in **b** and **c**, respectively. Scale bars, 15 μm. **C**, WISH for *pdgfaa*, *pdgfab* and *pdgfbb* at 24 hpf. Arrows point at signals at the ventral side of caudal somites. **D**, Quantification of pERK$^+$ cells in the tail of control, pdgfaa-, pdgfab- and pdgfbb-overexpressing embryos at 24 hpf (mean±SD; n=10, 16, 14 and 13 embryos for control, pdgfaa, pdgfab and pdgfbb, respectively, from a single experiment. **, P≤0.01; *, P<0.05; Student's *t*-test). **E**, Time-lapse confocal imaging from 22 to 26 hpf of *Tg(pax3a:eGFP; TCF:nls-mCherry)* embryos treated with 0.2 % DMSO (control) or 2 μM AS605240 from 20 hpf. White and blue arrows indicate migrating SCPs and neural crest cells, respectively.

Fig. S4

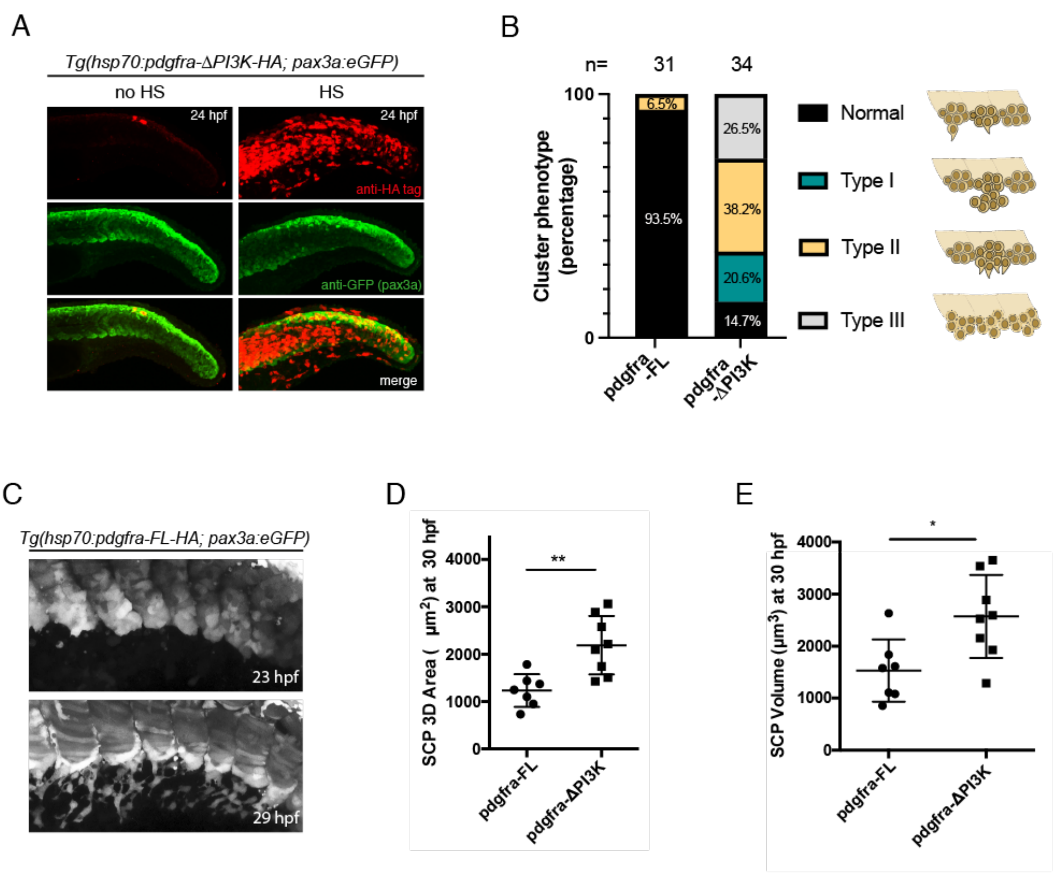


**Figure S4. Pdgfra-ΔPI3K induces defects in cluster integrity, emergence and migrating morphology of SCPs. A**, Immunofluorescence at 24 hpf for HA-tag (red) and pax3a:eGFP (green) in the tail of hsp70:pdgfra-ΔPI3K-HA injected embryos without heat-shock (no HS) and after heat-shock at 20 hpf (HS). **B,** Bar graph representing the frequency of each VC phenotype in Pdgfra-FL and Pdgfra-ΔPI3K expressing embryos. Typical phenotypes observed at 23 hpf in hsp70:pdgfra-ΔPI3K heat-shocked embryos are shown; Type **I**, 'delamination/overflow' phenotype; Type **II**, 'simultaneous migration' phenotype; Type **III**, 'loose cluster' phenotype. n=31 and 34 for pdgfra-FL and pdgfra-ΔPI3K embryos, from 10 independent experiments. **C**, Confocal maximum projections of a *Tg(hsp70:pdgfra-FL-HA; pax3a:eGFP)* embryo at 23 and 29 hpf. (**D**,**E)** 3D surface area (**D**) and volume (**E**) of individual SCP leader cells detached from their followers, for hsp70:pdgfra-FL and hsp70:pdgfra-ΔPI3K embryos at 30 hpf (n=7 and 8 cells for pdgfra-FL and pdgfra-ΔPI3K embryos, from two independent experiments; mean±SD; **, *P*=0.003; *, *P*=0.0143. Student's *t*-test)

Fig. S5

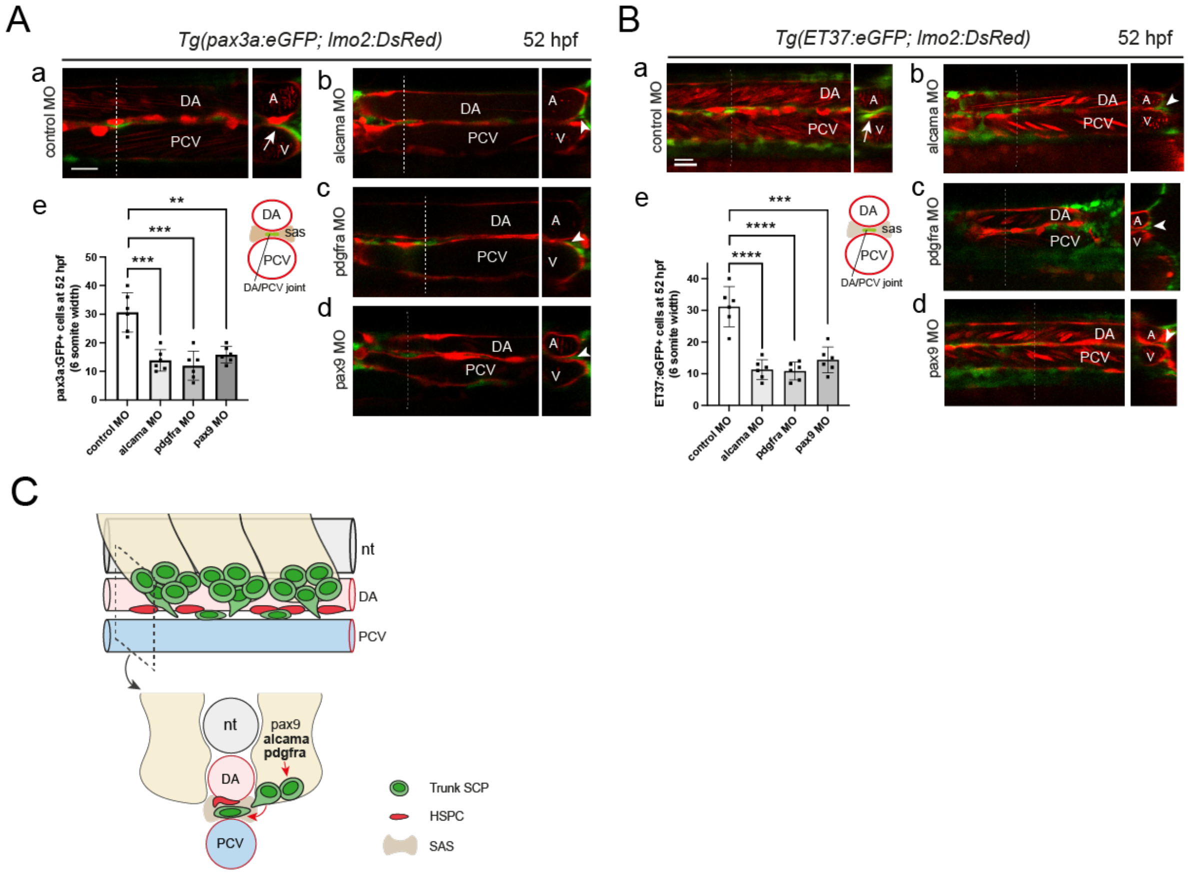


**Figure S5. Effect of alcama, pdgfra and pax9 MOs on stromal cell development in the trunk. A,B(a-d)** Confocal sections at 52 hpf of the trunk region of *Tg(pax3a:eGFP; lmo2:DsRed)* embryos (**A**) or *Tg(ET37:eGFP; lmo2:DsRed)* embryos (**B**) injected with control (**a**), alcama (**b**), pdgfra (**c**) and pax9 (**d**) MOs. Dashed vertical lines indicate the position where a corresponding optical transverse section is shown to the right. Arrows and arrowheads point at stromal cells located in the DA/PCV joint or more lateral to it (sub-aortic space), respectively. **A,B(e)**, Quantification of pax3a:GFP$^{+}$ (**A**) or ET37:eGFP$^{+}$ (**B**) cells in live embryos injected with control, alcama, pdgfra or pax9 MO at 36 hpf. Counting was performed over a 6-somites width. Mean±SD; n=6 embryos for each group; ****, $P \leq 0.0001$; ***, $P \leq 0.001$; **, $P \leq 0.01$; Student's *t*-test. DA or A, dorsal aorta; PCV or V, posterior cardinal vein.; sas, sub-aortic space. **C**, A schematic diagram of SCP development in the trunk region, showing that SCPs migrating from the somite VCs (sclerotome) position themselves just ventral to the dorsal aorta, in contact with the hemogenic endothelium and HSPCs that emerge from it via EHT, and/or with the dorsal side of the PCV, through which these HSPCs will enter circulation. The close presence of these somite-derived stromal cells appears essential for HSPC emergence.

Fig. S6

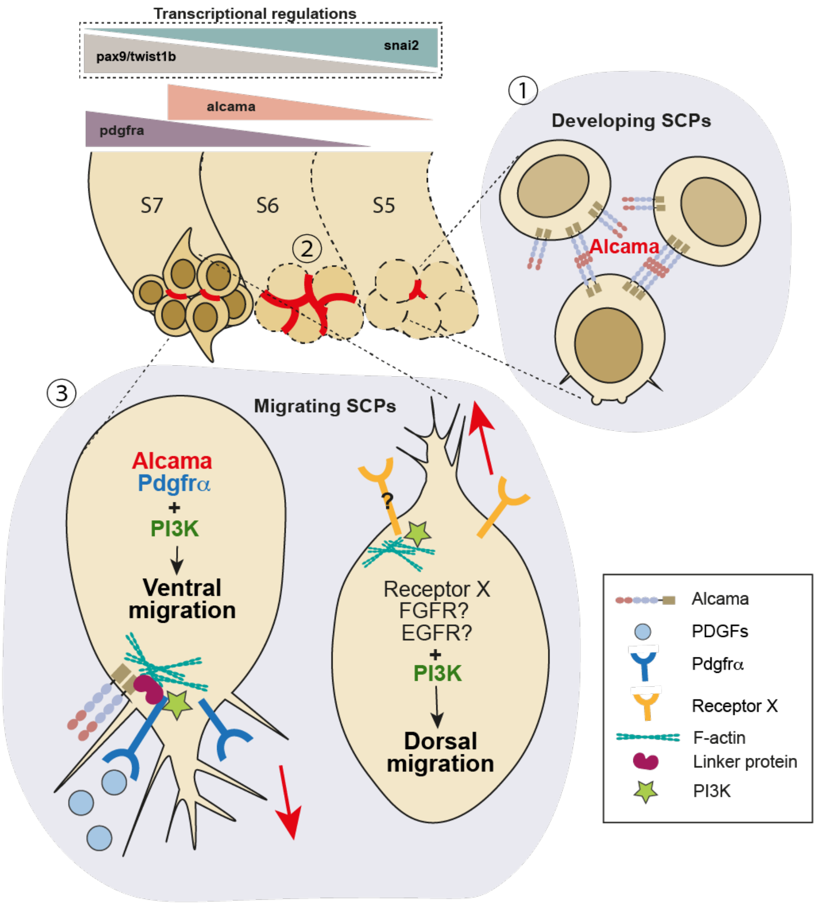


**Figure S6. Schematic representation of SCP emergence.** ① EMT occurs in the ventral part of caudal somites (around maturation stage S5), making the sclerotome cluster morphologically apparent. Alcama first appears at the center of the cluster. ② Alcama progressively spreads to all cell-cell interfaces within the cluster (around S6), ensuring selective adhesion among them. ③ By stage S7, sclerotome cells initiate semi-collective dorsal-ward migration, or ventral-ward migration to become SCPs. Alcama and Pdgfra are involved in the regulation of F-actin in SCPs through a molecular crosstalk, and ventral migration is triggered by PI3K, which is activated downstream of Pdgfra stimulated by PDGFs. PI3K is also involved in dorsal migration of sclerotome cells, but the upstream signals have not yet been identified. Among sclerotomal TFs, Snai2 activates *alcama* (likely indirectly) in younger somites, while Twist1b activates *pdgfra* expression in more developed somites. Considering that Pax9 shows inhibitory activity against *alcama* and *pdgfra*, and its expression level increases with somite maturation, it may exert its effect on cluster formation by balancing the activating effects of Snai2 and Twist1b.

#### Movie Legends

**Movie 1. VE-DIC imaging of ventral somite cells forming clusters.** The caudal region covering somite maturation stages S4 to S6 in a wild-type embryo was imaged every 10 sec. for 48 min. Arrows indicate the central lumen observed upon cluster formation. Scale bar, 20 µm.

**Movie 2. pax3a:eGFP$^{medium}$ expression marks somite VCs and SCPs emigrating from them.** 3D reconstitution from the time-lapse confocal imaging of a developing *Tg(pax3a:eGFP; TCF:nls-mCherry)* embryo imaged every 6 min for 10 hrs, starting at 24 hpf. Anterior to the left, dorsal to the top. pax3a:eGFP$^{medium}$ stromal progenitors emigrate in ventral direction from the ventral border of caudal somites. Some of these cells have inherited TCF:nls-mCherry$^{+}$ from their somitic origin. Pax3a:eGFP$^{high}$ neural crest cells that are pigment cell precursors can also be seen migrating, from the dorsal side of the spine over the medial side of the somites and then beginning to arrive in the CHT by the end of the sequence.

**Movie 3. Live imaging of an Alcama-FL-eGFP$^{+}$ SCP.** Following injection of a *Tg(cspg4:Gal4;UAS:RFP)* embryo with a UAS:alcama-FL-eGFP construct at the 1-cell stage, GFP$^{+}$ SCPs were imaged for 20 hours every 6 min. from 23 hpf. The left panel shows the overlay of RFP and GFP signals, and the intense GFP signal at cell contact points. The main GFP$^{+}$ cell followed up here undergoes mitosis at t=10h36, after which (by t=11h) intense GFP signals underline the interface of the two daughter cells. Scale bar, 20 µm.

**Movie 4. Live imaging of an Alcama-ΔN-eGFP$^{+}$ SCP.** Following injection of a *Tg(cspg4:Gal4;UAS:RFP)* embryo with a UAS:alcama-ΔN-eGFP construct at the 1-cell stage, a GFP$^{+}$ SCP was imaged for 16 hours every 6 min. from 23 hpf. The left panel shows the overlay of RFP and GFP signals, and the right panel the GFP signal only. Scale bar, 10 µm.

**Movie 5. Live imaging of an Alcama-ΔPDZ-eGFP$^{+}$ SCP.** Following injection of a *Tg(cspg4:Gal4;UAS:RFP)* embryo with a UAS:alcama-ΔPDZ-eGFP construct, a GFP$^{+}$

SCP was traced for 7 hours. Images were acquired at 6 min intervals from 25 hpf. The left panel shows the overlay of RFP and GFP signals, and the right panel the GFP signal only. Scale bar, 10 μm.

**Movie 6. Pdgfra deficiency affects the number and migration range of SCP-derived stromal cells, and thereby venous plexus structure.** Time-lapse confocal imaging of *Tg(pax3a:eGFP; lmo2:DsRed)* embryos injected with control or pdgfra MO. Images were acquired from 38 hpf for 20 hrs at 6 min intervals. $GFP^{high}$ cells forming continuous strings at the ventral and lateral border of somites belong to the dermomyotome (also called External Cell Layer), whereas $GFP^{high}$ cells migrating into the CHT are neural crest-derived pigment cells. SCP derived stromal cells are $GFP^{medium}$. The Lmo2:Dsred transgene highlights vascular cells, and more weakly red blood cells and HSPCs. Note that in the morphant embryo, the endothelial cells that formed the venous plexus did not migrate further ventral-wards than the SCP-derived stromal cells, and this led to a correspondingly narrower venous plexus, often reduced to a single convoluted tube. Anterior to the left. Scale bar, 30 μm.

**Movie 7. Pdgfra-ΔPI3K expressing SCPs show abnormal migration behavior and morphology.** Time-lapse confocal imaging of SCPs expressing pdgfra-FL-lifeact-GFP or pdgfra-ΔPI3K-lifeact-GFP. Images were acquired for 10 hrs at 6 min intervals from 24 hpf. $GFP^+$ SCPs were located in both cases at the ventral border of a somite VC at timepoint zero. Scale bar, 10 μm.

**Movie 8. Dynamics of pax3a:$GFP^+$ mesenchymal cells in the sub-aortic space, between DA and PCV**. Time-lapse confocal imaging of a *Tg(pax3a:eGFP; kdrl:ras-mCherry)* embryo. Images were acquired for 17 hrs at 6 min. intervals from 42 hpf. Scale bar, 50 μm.

**Supplementary information**

Table 1. Primers used for BAC transgenesis

| Primer | Sequence | Reference |
|---|---|---|
| pIndigobac_iTol2 F | TTCTCTGTTTTTGTCCGTGGAATGAACAATGGAAGTCC GAGCTCATCGCTCCCTGCTCGAGCCGGGCCCAAGTG | Suster et al.,2011 Bussmann and Schult-Merker, 2011 |
| pIndigoBAC_iTol2 R2 | AGCCCCGACACCCGCCAACACCCGCTGACGCGAACCC CTTGCGGCCGCATATTATGATCCTCTAGATCAGATC | Suster et al.,2011 Bussmann and Schult-Merker, 2011 |
| Cspg4_GAL4FF F | CTCTCCAGGTCCCAAAGTGGCCACAGAGACTCAGAGA CTCGGACTAAAGTgccaccatgAAGCTACTGTCTTCTATCG AAC | This study |
| Cspg4_frt-kan R | aggtataggagtgccaggaagagggcagacaggagcggacacggggctct CCGCGTGTAGGCTGGAGCTGCTTC | This study |

Table 2a. Primers used for the synthesis of WISH probes

| Primer | Sequence |
|---|---|
| alcama-WISH-F | CCTGCCGACGGTTATAGGTC |
| alcama-WISH-R | aagcttTAATACGACTCACTATAGGGGGCCGGTAATTCTTGGACCA |
| pdgfaa-WISH-F | TAGAAAGGCATGTTCCCCGG |
| pdgfaa-WISH-R | aagcttTAATACGACTCACTATAGGGAGAGTGATCCAAGAGCTGCG |
| pdgfbb-WISH-F | AAGAGCGGGGACAAAAGTGG |
| pdgfbb-WISH-R | aagcttTAATACGACTCACTATAGGGAAGAGCGGGGACAAAAGTGG |
| snai2-WISH-F | ACACTGAGAGGCCTGCATTC |
| snai2-WISH-R | aagcttTAATACGACTCACTATAGGGGGCATGTTCAAAACCTCAAACC |

Table 2b. Plasmids used for the synthesis of WISH probes

| Gene name | Construct | Reference |
|---|---|---|
| cspg4 | cspg4:pJC53.2 | Wang et al., 2014[1] |
| myb | cmyb:pBK-CMV | Thompson et al., 1998[2] |
| pax9 | pax9:pPICT3 | Kudo et al., 2004[3] |
| pdgfab | pdgfab:pExpress1 | This study |
| pdgfra | pdgfra:pCR4 | Eberhart et al., 2008[4] |
| twist1a | twist1a:pExpress1 | This study |
| twist1b | twist1b:pExpress1 | This study |

Table 3. Antibodies used in this study

| Antibody name | Dilution | Reference number, Manufacturer |
|---|---|---|
| zn-8 (anti-Alcama) | 1:100 | zn-8, DSHB |
| Phospho-p44/42 MAPK (ERK1/2) | 1:100 | #4370, Cell Signaling Technology |
| anti-HA.11 Epitope Tag | 1:50 | 901501, Biolegend |
| Chicken anti-GFP | 1:800 | ab13970, Abcam |
| Rabbit anti-DsRed | 1:300 | 632496, Takara |
| anti-Chicken-AlexaFluor 488 | 1:300 | A-11039, ThermoFisher Scientific |
| anti-Mouse-HRP | 1:300 | F-21453, ThermoFisher Scientific |
| anti-Rabbit-HRP | 1:300 | G-21234, ThermoFisher Scientific |

Table 4. Primers used for mutagenesis

*UAS:alcama-ΔN-eGFP*

| Primer | Sequence |
|---|---|
| G-UAS-alcama-SP F | GACGCGTGGATCCACCGGTCGCCACGCCACCATGCATTCGGTTATCTGCCTTTTCGGTG |
| G-UAS-alcama-SP R | GTAGACTCACCTTCTCAGTGGGCGGCAGGCAGCTCCCTGG |
| G-UAS-alcama-dN F | TGCTCCAGGGAGCTGCCTGCCGCCCACTGAGAAGGTGAGTCTACAG |
| G-UAS-alcama R | TGAACAGCTCCTCGCCCTTGCTCACGCCTGCTCCGACATCTGCTTTATGATTGTTCTCCTCC |

*UAS:alcama-ΔPDZ-eGFP*

| Primer | Sequence |
|---|---|
| G-UAS-alcama-SP F | GACGCGTGGATCCACCGGTCGCCACGCCACCATGCATTCGGTTATCTGCCTTTTCGGTG |
| G-UAS-alcama-dPDZ R1 | TCCGTCCGCCTGGCATTTCAGAGTCACATCATCAC |
| G-UAS-alcama-dPDZ F2 | TGATGATGTGACTCTGAAATGCCAGGCGGACGGAAAC |
| G-UAS-alcama-dPDZ R2 | TCC**C**GCTGCCTTGTCT**CACCA**TCTTGATATACAACCAGTAGATGAGTCCCACCAG |
| G-UAS-alcama-dPDZ F3 | ATCAAGA**TGGTG**AGACAAGGCAGC**G**GGAAGACCGGAGAGAAGGAGAC |
| G-UAS-alcama R | TGAACAGCTCCTCGCCCTTGCTCACGCCTGCTCCGACATCTGCTTTATGATTGTTCTCCTCC |

*UAS:alcama-FL-eGFP*

| Primer | Sequence |
|---|---|
| G-UAS-alcama-SP F | GACGCGTGGATCCACCGGTCGCCACGCCACCATGCATTCGGTTATCTGCCTTTTCGGTG |
| G-UAS-alcama-dPDZ R1 | TCCGTCCGCCTGGCATTTCAGAGTCACATCATCAC |
| G-UAS-alcama-dPDZ F2 | TGATGATGTGACTCTGAAATGCCAGGCGGACGGAAAC |
| G-UAS-alcama R | TGAACAGCTCCTCGCCCTTGCTCACGCCTGCTCCGACATCTGCTTTATGATTGTTCTCCTCC |

*hsp70:pdgfra-ΔPI3K-HA*

| Primer | Sequence |
|---|---|
| G-hsp70-pdgfra-dn F1-2 | CCTGGAATTCGGTACCCTCGAGGATATCGCCACCATGTTCCCGGTGCTGCCACAGTCAGTTCAGGCTC |
| G-hsp70-pdgfra-dn R1 | AGGTTCAACACACCCTTGTAACTGGTCTCGCCGATTTTGAGGAGGTTGGTGCTG |
| G-hsp70-pdgfra-dn F2 | GCACCAACCTCCTCAAAATCGGCGAGACCAGTTACAAGGGTGTGTTGAACCTG |
| G-hsp70-pdgfra-dn R2 | TGGGGACG**A**ACTGCATGGTGTCTGCCTGCTTCATGTCCATA**A**AATCACCTTTCCCCTC |
| G-hsp70-pdgfra-dn F3 | AGGTGATT**T**TATGGACATGAAGCAGGCAGACACCATGCAGT**T**CGTCCCCATGCTGGAAATG |
| G-hsp70-pdgfra-dn R3 HA-tag | GTATGGCTGATTATGATCGCGGCCGCGGATCCTCATGCGTAATCAGGCACATCATAAGGATAAGCGTAGTCTGGGACGTCGTATGGGTAGCCTGCTCCCAGGAAGCTGTCCTCCACCAGGTC |

*hsp70:pdgfra-FL-HA*

| Primer | Sequence |
|---|---|
| G-hsp70-pdgfra-dn F1-2 | CCTGGAATTCGGTACCCTCGAGGATATCGCCACCATGTTCCCGGTGCTGCCACAGTCAGTTCAGGCTC |
| G-hsp70-pdgfra-dn R1 | AGGTTCAACACACCCTTGTAACTGGTCTCGCCGATTTTGAGGAGGTTGGTGCTG |
| G-hsp70-pdgfra-dn F2 | GCACCAACCTCCTCAAAATCGGCGAGACCAGTTACAAGGGTGTGTTGAACCTG |

| | |
|---|---|
| G-pdgfra-wt R1 | TGGGGACGTACTGCATGGTGTCTGCCTGCTTCATGTCCATATAATCAC |
| G-pdgfra-wt F1 | AGGTGATTATATGGACATGAAGCAGGCAGACACCATGCAGTACGTC |
| G-hsp70-pdgfra-dn R3 HA-tag | GTATGGCTGATTATGATCGCGGCCGCGGATCCTCATGCGTAATCAGGCACATCATAAGGATAAGCGTAGTCTGGGACGTCGTATGGGTAGCCTGCTCCCAGGAAGCTGTCCTCCACCAGGTC |

Table 5. Primers used  for cloning

| Primer | Sequence |
|---|---|
| Alcama_SC_frg1 F1 | GCCGGACTGTATAAAGGAGAACC |
| Alcama_SC_frg1 R1 | AAGCTCACTGTGACGATCTGAG |
| Alcama_SC_frg2 F2 | AAGGGAAAGAAGGTCACGGTG |
| Alcama_SC_frg2 R2 | AGTGTGACAAGGCCTCTTTCC |
| Pdgfaa_SC F1 | CTGCGCTGGGACACTTTTG |
| Pdgfaa_SC R1 | AGAGTGATCCAAGAGCTGCG |
| Pdgfab_SC F1 | GGTGCATCGGGTCATTTATAG |
| Pdgfab_SC R1 | ATGTGGTTTCACCTCATGTCC |
| Pdgfbb_SC F1 | ATATTTGCTCGCGTTAAGTGG |
| Pdgfbb_SC R1 | GATTGCATCCCTCTGAACATC |
| Twist1b_SC F1 | ACCCTCATGCTGGAATAACG |
| Twist1b_SC R1 | TCCTCGTGTTTTCCCAGCTC |
| G-pax9-ORF5' F | ACACTATAGAACAAGTTTCGGTCCGGAATTCGTTACCATGGAGCCAGCCTTTGGGGAG |
| G-pax9-ORF3' R | TACGACTCACTATAGGGACCACTCCTCGAGTCACAGCTGTGGGGAGAGAGAGC |
| G-snai2-ORF5' F | ACTATAGAACAAGTTTCGGTCCGGAATTCGTTACCATGCCTCGTTCATTCCTAGTAAAG |
| G-snai2-ORF3' R | ATACGACTCACTATAGGGACCACTCCTCGAGTCAGTGTGCGATGCAACAGCCAG |

Table 6. MOs used in this study

| MO name | MO sequence (5' -> 3') | reference |
|---|---|---|
| MO1-alcama | GTCCGGCGACAGTCTCAATAGAGAG | Choe et al., 2013 |
| MO1-pax9 | CCAAAGGCTGGCTCTAGTTATGCAG | Swartz et al., 2011<br>Charbord et al., 2014 |
| MO2-pdgfra | TTCGAGACATCTGCAAGGAGATATT | French et al., 2014 |
| MO1-snai2 | ATACATGTCATTTTCTCACCCGTGT | Charbord et al., 2014<br>Bickers et al., 2018 |
| MO1-twist1a | ACCTCTGGAAAAGCTCAGATTGCGG | Das et al., 2012 |
| MO3-twist1b | TTAAGTCTCTGCTGAAAGCGCGTG | Das et al., 2012 |

Table 7. Primers used for qPCR

| Primers | Sequence |
|---|---|
| alcama-qPCR-F | AGTGGAGTGTCAACGGAACC |
| alcama-qPCR-R | CGAGTTTATTGGTCACGAGGC |
| pax9-qPCR-F | CCACTCTTCCTGGACATATGGC |
| pax9-qPCR-R | TTCTAGTTTGGCGCTGGGAAAG |
| pdgfra-qPCR-F | ACGCTGAGTGATGTCTGGTC |
| pdgfra-qPCR-R | AGGTTTGGTCATTCGGTATCC |

| | |
|---|---|
| snai2-qPCR-F | TGCAGGGACACATTAGAACAC |
| snai2-qPCR-R | TGCACTGGTATTTCTTCACGTC |
| twist1a-qPCR-F | TGTCAACATCCCACTAACGCAC |
| twist1a-qPCR-R | TGACGCTCCAGAATTTTCCC |
| twist1b-qPCR-F | ACACAAAGTTGCTTGGAATACC |
| twist1b-qPCR-R | CTCGCTTAAGTCTCTGCTG |

Table 8. Primers used for the cloning of alcama and pdgfra promoters

| **Primer** | **Sequence** |
|---|---|
| G alcama promoter F1-2 | TCTTACGCGTGCTAGCCCGGGCTCGAGCAGCATTGATGGTTCCATGAAGA |
| G alcama promoter R1-2 | GAAACCAACGTAGGACCTTAAGACCCTCCAGGTAAGAACATG |
| G alcama promoter F2 | GACATGTTCTTACCTGGAGGGTCTTAAGGTCCTACGTTGGTTTC |
| G alcama promoter R2-2 | GCTGCAACCTATCACTGGACAACATCCATACACTTTAACACAAAATATTTAAC |
| G alcama promoter F3-2 | GTGTTAAAGTGTATGGATGTTGTCCAGTGATAGGTTGCAGC |
| G alcama promoter R3-2 | CTTTACCAACAGTACCGGAATGCCAAGCTTATTGAGAGTTGAGTGCGCTG |
| G pdgfra promoter F1 | TTACGCGTGCTAGCCCGGGCTCGAGTCACAATATATCTGTTTCATAAGG |
| G pdgfra promoter R1 | CATCGTCTAACCACTCTTATAAAGTCGTTAACAACATTATAAGTAGC |
| G pdgfra promoter F2 | CAAGCTACTTATAATGTTGTTAACGACTTTATAAGAGTGGTTAGAC |
| G pdgfra promoter R2 | CATGGCAGGCTGCAGATAATGGCTCATTATGAATTAGTACCCTACG |
| G pdgfra promoter F3 | GTCGCGTAGGGTACTAATTCATAATGAGCCATTATCTGCAGCCTG |
| G pdgfra promoter R3 | CTTTACCAACAGTACCGGAATGCCAAGCTTAGCTTAATGCTACAGTCCATC |

Table 9. Injection conditions for Luciferase assay (amount per egg)

| **pGL-alcama-luc** | **pRL-TK** | **pax9 MO** | **pax9 mRNA** | **snai2 MO** | **snai2 mRNA** |
|---|---|---|---|---|---|
| 15 pg | 1.5 pg | 6 ng | - | - | - |
| | | - | 50 pg | - | - |
| | | - | - | 8 ng | - |
| | | - | - | - | 100 pg |

| **pGL-pdgfra-luc** | **pRL-TK** | **snai2 MO** | **snai2 mRNA** | **twist1a+1b MO** | **twist1b mRNA** |
|---|---|---|---|---|---|
| 15 pg | 1.5 pg | 8 ng | - | - | - |
| | | - | 100 pg | - | - |
| | | - | - | 2 ng each | - |
| | | - | - | - | 100 pg |
